# Coordinated plant and microbial transcriptional responses to oil-sands process-affected water

**DOI:** 10.64898/2026.01.14.699257

**Authors:** J.E Nweze, S. Morvan, A. Samad, M-J. Bergeron, D. Degenhardt, J. Tremblay, K. Symonds, D.G. Muench, C. Martineau, E. Yergeau

## Abstract

Constructed wetland treatment systems (CWTSs) are promising options for treating oil-sands process-affected water (OSPW), which contains toxic naphthenic acid fraction compounds (NAFCs). However, the molecular mechanisms underlying NAFCs attenuation by plants and root microbes remain poorly resolved. In our previous mesocosm study, common cattail (*Typha latifolia*) increased NAFCs removal by 2.5-fold relative to unplanted controls with no significant effect on plant growth. Here, using RNA from the same experimental system, we applied metatranscriptomics to 40 root samples collected over 60 days to examine plant and active-microbial responses to OSPW exposure. The active-root-associated-microbial community was dominated by *Pseudomonadota*, which showed a slight increase with exposure to OSPW. *Burkholderiales* were the most active family, though their relative activity decreased in OSPW systems, where *Flavobacteriaceae* (*Bacteroidota*) activity increased. Clear microbial-community shifts were driven by time and OSPW exposure. Although 18 previously proposed microbial NAFC-degradation genes were not differentially expressed, 42 other genes with potential roles in NAFC or related organic compound degradation showed differential expression in OSPW-filled mesocosms. This activity was dominated by specific oxidoreductases from *Burkholderiales* and *Rhizobiales*. Crucially, host plant actively responded to OSPW, robustly up-regulating genes encoding oxidoreductases, transporters, and glycosyltransferases, some of which are related to xenobiotic stress and detoxification. Taken together, these results show coordinated plant and microbial transcriptional responses in a system where NAFC removal had already been measured chemically. They help explain the response of OSPW-exposed mesocosms, but the observed patterns likely reflect the broader OSPW mixture rather than NAFCs alone.

## Introduction

Oil sands process-affected water (OSPW), generated during the hot-water extraction process of bitumen, contains a complex mixture of dissolved organic and inorganic compounds, with naphthenic acid fraction compound (NAFCs) being one of the major toxic pollutants [1]. NAFCs are a large and diverse group of organic acids, including classical naphthenic acids, naturally-occurring dissolved organic matter and any bitumen-derived components [2]. Several treatment technologies proposed for the remediation of OSPW are energy-intensive, costly, and require *ex-situ* treatments [3]. This highlights the need for sustainable *in situ* alternatives, including plant- and microbe-based systems. However, the actively expressed microbial pathways, functional genes, and plant-responses associated with NAFC attenuation in operating wetland treatment systems remain poorly understood [4].

Microorganisms are central to bioremediation because of their remarkable ability to break down complex organic pollutants through a wide range of enzymatic processes, ultimately leading to degradation, detoxification, or immobilization [5]. Both bacteria and fungi can metabolize toxic hydrocarbons under aerobic or anaerobic conditions, using them as sources of carbon and energy and producing less harmful by-products in the process [6]. This ability is linked to a range of enzyme systems, including oxidoreductases (e.g., dehydrogenases, cytochrome P450s), hydrolases, laccases, dehalogenases, lipases, and proteases, involved in the breakdown of hydrocarbons in various environments [7]. Although microbial degradation of NAFCs and NAFC-like compounds has been reported [4], only a few NAFC-degradation microbes, genes, and pathways have been identified.

Beta-oxidation (ring cleavage) with α-oxidation, ω-oxidation, aromatization or CoA-thioester activation has been identified as potential pathways for aerobic NAFC degradation, and benzoyl-CoA with nitrate, sulfate, methanogens or iron reducers as potential anaerobic NAFC degradation pathways [4]. Yet, most prior studies investigating the role of microbial communities in NAFC degradation have restricted their analyses to community-level profiling (e.g., DGGE, 16S rRNA sequencing) in tailings ponds or upland and wetland systems [8,9], leaving critical gaps about gene expression or directly linking microbes to active metabolic processes.

Complementing the capabilities of microbes, plants also play an important role in bioremediation by using different phyto-mechanisms to uptake, detoxify, immobilize, stabilize, or degrade diverse organic pollutants [10]. Their roots, particularly in constructed wetland treatment systems (CWTSs), enhance microbial degradation by providing root exudates and oxygen, and restructuring the surrounding habitat, and stimulate beneficial microbial activity [11]. This mutualistic interaction results in both sequestration and partial detoxification of pollutants, as observed in wetland plants like *Typha* spp. (cattails), which are capable of uptaking and transforming heavy metals and organic compounds within their tissues while fostering diverse root-associated microbial communities [10,12]. In contrast to microbes, there is limited knowledge on plant NAFC-degradation and transport genes [13].

In our previous study, *Typha latifolia* planted in greenhouse CWTS mesocosms significantly reduced total NAFCs relative to unplanted controls and also altered water and substrate chemistry [14]. Building on that system, we analysed RNA extracted from *T. latifolia* roots harvested from the same mesocosms to characterize the active microbial community and host transcriptional response. The aim of the present study was not to remeasure NAFC removal, but to interpret plant and microbial transcriptional responses in the context of previously quantified attenuation. Specifically, we compared root metatranscriptomes from OSPW- and reverse osmosis water (ROW)-filled mesocosms over 60 days to identify active taxa and differentially expressed plant and microbial genes associated with exposure to OSPW.

## Materials and methods

### Study site, materials, plant propagation and mesocosm setup

The source of coarse sand tailings (CST) and oil sands process-affected water (OSPW), plant propagation of *Typha latifolia*, and the full greenhouse mesocosms design were described previously in Balaberda *et al*. [14]. Briefly, the experiment was conducted under controlled greenhouse conditions to minimize seasonal variability rather than to evaluate seasonal effects. Common cattail (*Typha latifolia*) was propagated from field-collected seeds (zone DM1.3; dry mixedwood) for 3 months in peat-filled styroblock containers before transplanting. Twelve plants were transplanted into each mesocosm (average height of 120.8±17.0 cm). Each contained a 20 cm layer of CST (85.6 L) and either OSPW or reverse osmosis water (ROW; 106.9 L) maintained 25 cm above the substrate. For the metatranscriptomic component, we used a 2×2 factorial design with four replicates per treatment (n=16): ROW without plants, ROW planted with *T. latifolia*, OSPW without plants, and OSPW planted with *T. latifolia*. Mesocosms were acclimated with ROW before a 24-hour flush with the assigned water type and the start of the 60-day exposure period. Root sampling was destructive and performed at day 0, 6, 19, 40, and 60, yielding 40 root samples for sequencing.

Both CST and OSPW were chemically characterized in the paired mesocosm study from which the RNA for this work was obtained [14]. CST consisted predominantly of sand (98.0% ± 0.0), with minor silt and clay fractions. Relative to ROW, OSPW was characterized by higher salinity, alkalinity, and major ion concentrations typical of tailings pond water, including elevated Ca, Na, K, Cl, and SO _4_. In that same study, planted mesocosms altered water chemistry through reductions in salinity, alkalinity, and selected ions over time. Those measurements provide the chemical context for interpreting the transcriptional responses reported here.

### Root sampling and RNA extraction

Root samples were manually collected right after draining the ROW from the mesocosms at the end of the acclimation period (day 0) and 6, 19, 40 and 60 days after refilling the mesocosms with either ROW or OSPW (from this point referred to as D0, D6, D19, D40 and D60). Roots were immediately cleaned after sampling, snap-frozen in a dry ice-ethanol bath, shipped on dry ice, and stored at –80^°^C. Prior to RNA isolation, frozen root tissues (150-200 mg) were ground using 5 mm stainless steel beads and a TissueLyser II (Qiagen). RNA was extracted using the NucleoSpin RNA plant and fungi kit (Macherey-Nagel). ERCC ExFold RNA Spike-In Mix 1 (Invitrogen) was used as internal control and mixed together with PFL and PFR buffers. Two DNase treatments were performed, using the TURBO DNA-free kit (Invitrogen). Concentrated RNA extracts were recovered using the RNA Clean & Concentrator-25 kit (Zymo Research). Absence of residual DNA in the RNA extracts was confirmed by 16S PCR [primers 515F-Y: 5’-GTGYCAGCMGCCGCGGTAA [15] and 926R: 5’-CCGYCAATTYMTTTRAGTTT [16]. RNA quality and quantity were assessed using the 2100 Bioanalyzer (Agilent) and Qubit 3 (Invitrogen) instruments, respectively.

### Library preparation and sequencing

Ribosomal RNA (rRNA) was depleted from 2 µg of root RNA using the Pan-Plant riboPOOL kit (siTOOLs Biotech) and streptavidin-coated magnetic beads (New England Biolabs). Size-selective purification and concentration were performed using the SPRIselect bead-based reagent (Beckman Coulter) and the RNA Clean & Concentrator-5 kit (Zymo Research), respectively. RNA was eluted in 10 µl of water and quantified using the Qubit RNA High Sensitivity assay kit (Invitrogen). Libraries were prepared using the NEBNext Ultra II RNA Library Prep kit for Illumina (New England Biolabs). Based on the RNA insert size, fragmentation incubation time was set to 8 minutes at 94°C. NEBNext Multiplex Oligos were used for the PCR enrichment of adaptor ligated DNA step and the number of PCR cycles was set to eight, based on the rRNA depleted RNA input amount. Library size and concentration were assessed using the High Sensitivity DNA kit (Agilent) and 2100 Bioanalyzer and the Qubit dsDNA High Sensitivity assay kit (Invitrogen), respectively. Equimolar libraries (3 ng/µl) were pooled and sequenced at the Centre d’expertize et de services, Génome Québec (Montréal, Canada), using a 25B flow cell and the NovaSeq X Plus system (Illumina) for 2×100 cycles (paired-end mode). A phiX library was used as a control and mixed at 1% level.

### Microbial community and beta diversity

The microbial community in metatranscriptomic raw reads was profiled using SingleM v0.18.3 [17], which targets active 35 bacterial and 37 archaeal single-copy marker genes. The resulting OTU table was transformed into a matrix in R. Alpha diversity (Observed richness and Shannon index) was calculated on raw counts to assess community complexity over water type and time. We fitted linear mixed-effects models with water type, time, and their interaction as fixed effects, and included mesocosm as a random intercept to account for repeated sampling within the experiment. Beta diversity was evaluated using Bray-Curtis dissimilarity matrices calculated from relative abundance-transformed data, and visualized via Principal Coordinates Analysis (PCoA). Formal multivariate testing was conducted using PERMANOVA (adonis2, 999 permutations) with the model Bray-Curtis ∼ water type * time. Beta-dispersion was also tested with betadisper and permutation tests for both water type alone and the combined water-type-by-time grouping structure. Within SingleM, hierarchical clustering was performed using the UPGMA algorithm with jackknife support values [18]. The taxon-based dissimilarity between the samples across the time points was generated, and it ranges from 0.0 (low dissimilarity) to 1.0 (high dissimilarity).

### Metatranscriptomic read processing and assembly

Sequences were processed using an in-house bioinformatics pipeline [19] built on the GenPipes workflow [20]. Reads were quality-controlled using Trimmomatic v0.39 [21] and BBDUK (BBTools v38.1), and co-assembled using MEGAHIT v1.2.9 [22]. The QC-passed reads were mapped with BWA mem v0.7.17 [23] against contigs to assess the quality of the assembly. Reads were then mapped using STAR v2.7.11b [24] against the *T. latifolia* reference genome (NCBI: JAAWWQ000000000.1) to separate plant (mapped) from non-plant/microbial (unmapped) transcripts.

### Differential gene expression analysis

Transcript abundances were analysed using DESeq2 [25] to identify differentially expressed genes (DEGs) in cattail roots between OSPW and ROW at the same five time points (0, 6, 19, 40, and 60 days). Here, day 0 represents samples collected immediately after draining the ROW at the end of the acclimation period, before exposure to OSPW or fresh ROW treatments. Before running the analysis, genes with ≥10 counts in ≥2 replicates were retained. Benjamini-Hochberg [26] controlled false discovery rate (FDR), and genes were considered differentially expressed if the adjusted p-value was ≤0.05 and the log2-fold-change (log_2_FC) was ≥1.

### Gene prediction and assignment

Gene prediction used Prodigal v2.6.3 [27], and predicted genes were assigned to KEGG orthologs (KO) using HMMER v3.3.2 [28] against the KOFAM database [29] with a significance threshold of 10^−10^. Targeted homology searches for NAFC-degradation genes (e-value 1e^−5^) included: (1) 18 previously associated genes (e-value of 1e^−5^; [3]); (2) hydrocarbon degradation markers (cutoffs: ≥25% coverage, ≥30% identity, e-value ≤1e^−5^) using CANT-HYD database containing 37 HMMs [30] and HADEG database containing 259 proteins [31]. Functional profiling for biogeochemical cycles employed curated databases for methane (MCycDB), nitrogen (NcycDB), sulfur (SCycDB), and phosphorus (PCycDB). BLAST identity thresholds were ≥75% for McycDB [32], NcycDB [33], ScycDB **[34]** cyclings and 30% for PCycDB cycling **[35]**. Gene abundance profiles were generated by mapping QC-passed reads against predicted genes using BWA mem v0.7.17 **[23]**. Taxonomic assignment used BLAST (e-value = 10^−10^) against NCBI databases [36]. All the generated data were visualized in R **[37]**, using ggplot2 **[38]** and iTOL [39]. The generated scalable vector graphics (SVG) were annotated in Inkscape and exported as Portable Network Graphics (PNG).

## Results and Discussion

### Experimental baseline and NAFC removal

As context for the metatranscriptomic analysis, we first considered the plant performance and chemical outcomes previously measured in this same mesocosm experiment. *Typha latifolia* maintained >95% survival in both oil sands process-affected water (OSPW) and reverse osmosis water (ROW) **[14]**, confirming its resilience under the conditions used here **[40]**. Although some shoots showed 70-99% necrotic tissue, that visual damage was attributed in the paired study to an uncontrolled aphid infestation rather than to a significant OSPW effect on plant height, biomass, or overall vigour (p > 0.05; Fig. 1a). In the same experiment, planted OSPW mesocosms achieved 40% NAFC removal, compared with 16% in unplanted controls (Fig. 1b), and plant presence also influenced salinity, alkalinity, and ion concentrations in the system. We use that previously quantified removal as the functional backdrop for the metatranscriptomic patterns described below.

**Fig. 1.**
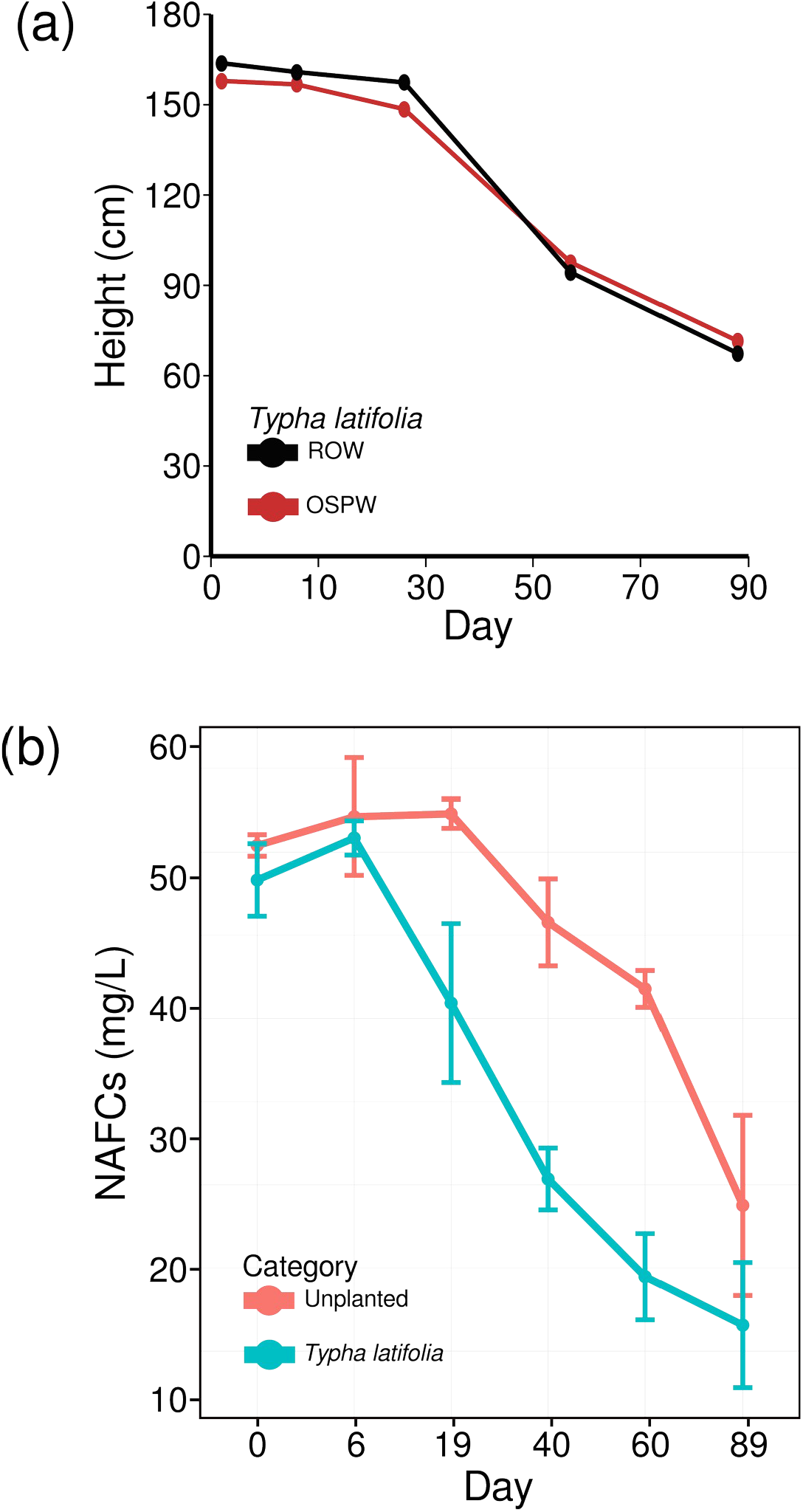
Plant performance and naphthenic acid fraction compound (NAFC) removal in the paired mesocosm study from which RNA was obtained for the present metatranscriptomic analysis. (a) Typha latifolia height over time in mesocosms supplied with oil sands process-affected water (OSPW) and reverse osmosis water (ROW). (b) Removal efficiency of NAFCs by planted and unplanted mesocosms supplied with OSPW. Time 0 indicates the sampling point before OSPW addition; Time 6-60 represent sampling days after adding OSPW or refilling with ROW. This figure is reproduced from Balaberda et al. [14] to provide the chemical and plant-performance context for the transcriptomic analyses presented here.

### *De novo* assembly information of the root microbial metatranscriptome

From our previous experimental setup (Table S1), RNA was extracted and sequenced from a total of 40 root samples of *Typha latifolia* grown in OSPW or ROW (Table S2A). After quality-filtering, the reads from all samples were assembled resulting in 17,541 contigs (Table S2B). The minimum and maximum contig lengths were 1000 and 16,907 (bp), respectively. Twenty-one of the contigs were greater than 10kb and the N50 was >= 1,604bp.

### Active microbial community in the root samples

We examined the composition and activity of the root-associated microbial community over time. Metatranscriptome profiling of root-associated prokaryotic communities in both OSPW and ROW mesocosms showed a clear dominance of *Pseudomonadota* among active microorganisms across all time points (Fig. 2a; Table S3 and Table S4A), peaking on D6 at 85% in OSPW and 87% in ROW after draining and refilling of water samples. After D6, the phylum declined but remained more abundant in OSPW than in ROW. As *Pseudomonadota* abundance reduced, the presence of *Bacteroidota*, the second most abundant phylum, increased. In OSPW samples, active *Bacteroidota* ranged from 11% to 18%, while in ROW it ranged from 8% to 18%. Its abundance increased steadily from D6 to D40 in both water types, followed by a decrease at D60. *Planctomycetota* was the third most abundant phylum in OSPW samples, ranging from 1% to 4%, except on D40, where *Actinomycetota* showed a slight increase at 1%. Interestingly, in our DNA-based study [13], while *Pseudomonadota* and *Bacteroidota* were also the most abundant phyla detected i*n T. latifolia* roots, *Actinomycetota* and *Verrucomicrobiota* were more abundant than *Planctomycetota*, indicating that this phylum might be less abundant but more active in roots exposed to OSPW.

**Fig. 2.**
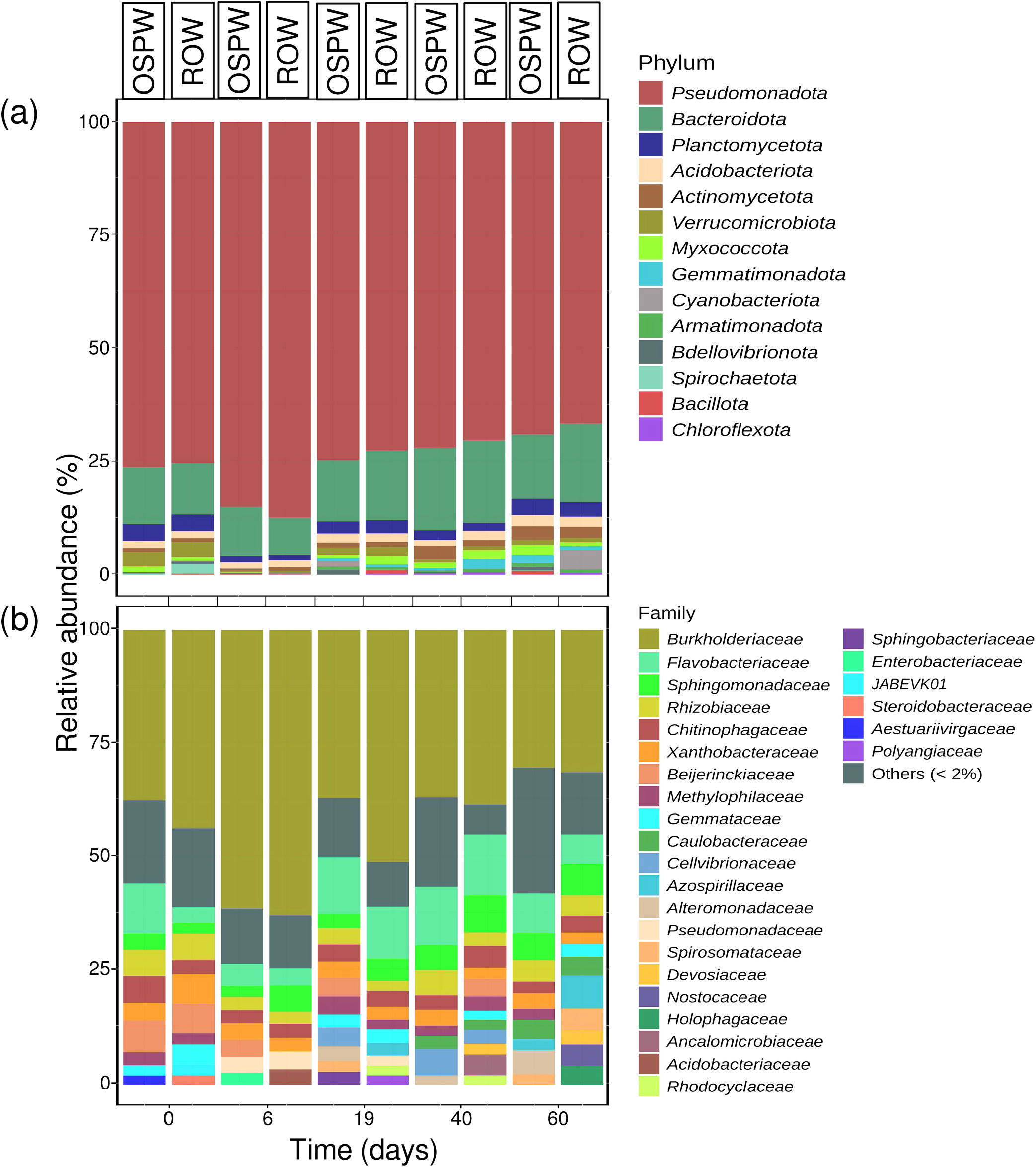
Active microbial profile of root-associated Typha latifolia in mesocosms containing oil sands process-affected water (OSPW) or reverse osmosis water (ROW). (a) Phylum-level relative abundance. (b) Family-level relative abundance. Profiles were generated from metatranscriptomic reads using the expressed universal-single-copy genes. Time 0 is before addition of OSPW; 6-60 represent sampling days after adding OSPW or refilling with ROW.

In the ROW samples, different phyla occupied the third position in relative abundance over time, including *Planctomycetota, Acidobacteriota, Gemmatimonadota*, and *Cyanobacteriota*. The dominance of *Pseudomonadota* in active cattail root communities is notable because many members of this phylum are associated with the metabolism of polycyclic aromatic hydrocarbons and other complex organic contaminants [43], and their dominance in active cattail root bacterial communities further supports the relevance of this plant species in remediation systems. However, in the present study this pattern should be interpreted as evidence of a community well suited to chemically complex conditions, rather than as direct proof of NAFC degradation. The trend is consistent with other *Typha*-planted tailings environments [44], although it differs from some natural wetlands, where *Actinobacteria, Bacillota, Pseudomonadota*, and *Planctomycetota* often co-dominate the root zone [45].

At the family level, *Burkholderiaceae* was consistently the most active at all time points in both water types (Fig. 2b; Table S3 and Table S4B), rising sharply at D6 (61% in OSPW; 63% in ROW), following draining and refilling with either OSPW or ROW, and then declining toward D60 (30% in OSPW; 31% in ROW). The abundance was higher in ROW than OSPW, driven primarily by increases in other families, especially *Flavobacteriaceae* (*Bacteroidota*). The same *Burkholderiaceae* family dominated our DNA-based study of *T. latifolia* roots, but it was followed by *Chitinophagaceae, Xanthobacteraceae, and Sphingomonadaceae*, in contrast to *J. balticus* roots, which were dominated by *Gallionellaceae [14]*. The dominance of *Burkholderiaceae* is consistent with studies that link the family’s abundance to tolerance of heavy metals and hydrocarbon contaminants [46] and its status as the predominant order in tailings layers [47]. Furthermore, the genus *Burkholderia* is widely known as a plant growth-promoting rhizobacteria (PGPR) that enhances plant resilience in stressful conditions [48]. *Flavobacteriaceae*, known to contribute positively to plant health, growth, and tolerance to abiotic stress [49], was the second most abundant family in OSPW (5–13%) and D19 (12%) and D40 (13%) in ROW, while other second-tier families—including *Beijerinckiaceae, Sphingomonadaceae Azospirillaceae* and *Xanthobacteraceae*— varied by time point in both water types. These families are root-associated taxa that support plant growth and resilience, even in *Typha* spp. [50,51].

### Microbial diversity between samples at different time points

Alpha diversity (Richness and Shannon) was comparatively stable between water types (Fig. 3a; Table S5A). Richness did not significantly differ between the water type (F = 1.51, p = 0.265), time (F = 1.09, p = 0.383), or their interaction (F = 0.90, p = 0.478), in contrast to patterns observed in metagenomes from the same sample [14]. Similarly, Shannon diversity showed no significant effect on the water type (F = 0.76, p = 0.418), time (F = 1.18, p = 0.344), or their interaction (F = 0.73, p = 0.583), consistent with metagenomic results [14]. These findings indicate that overall community diversity remained stable across treatments, although this does not preclude compositional or functional shifts. Beta diversity showed a clear overall multivariate signal across the experiment (Fig. 3b; Table S5B), indicating that bacterial community composition was reorganized over time and treatment. Bray–Curtis PCoA revealed structuring of samples by both time and water type, with replicate centroids shifting across sampling days. Separation between OSPW and ROW treatments in the hierarchical clustering was also evident, supporting an effect of water type on community composition (Fig. 3c). Samples from early and late time points formed distinct clusters, indicating that time was a major driver of community restructuring. As D0 samples were collected immediately after the acclimation phase and before the treatment period began, we interpret D0 primarily as a baseline reference rather than as a treatment-defined state.

**Fig. 3.**
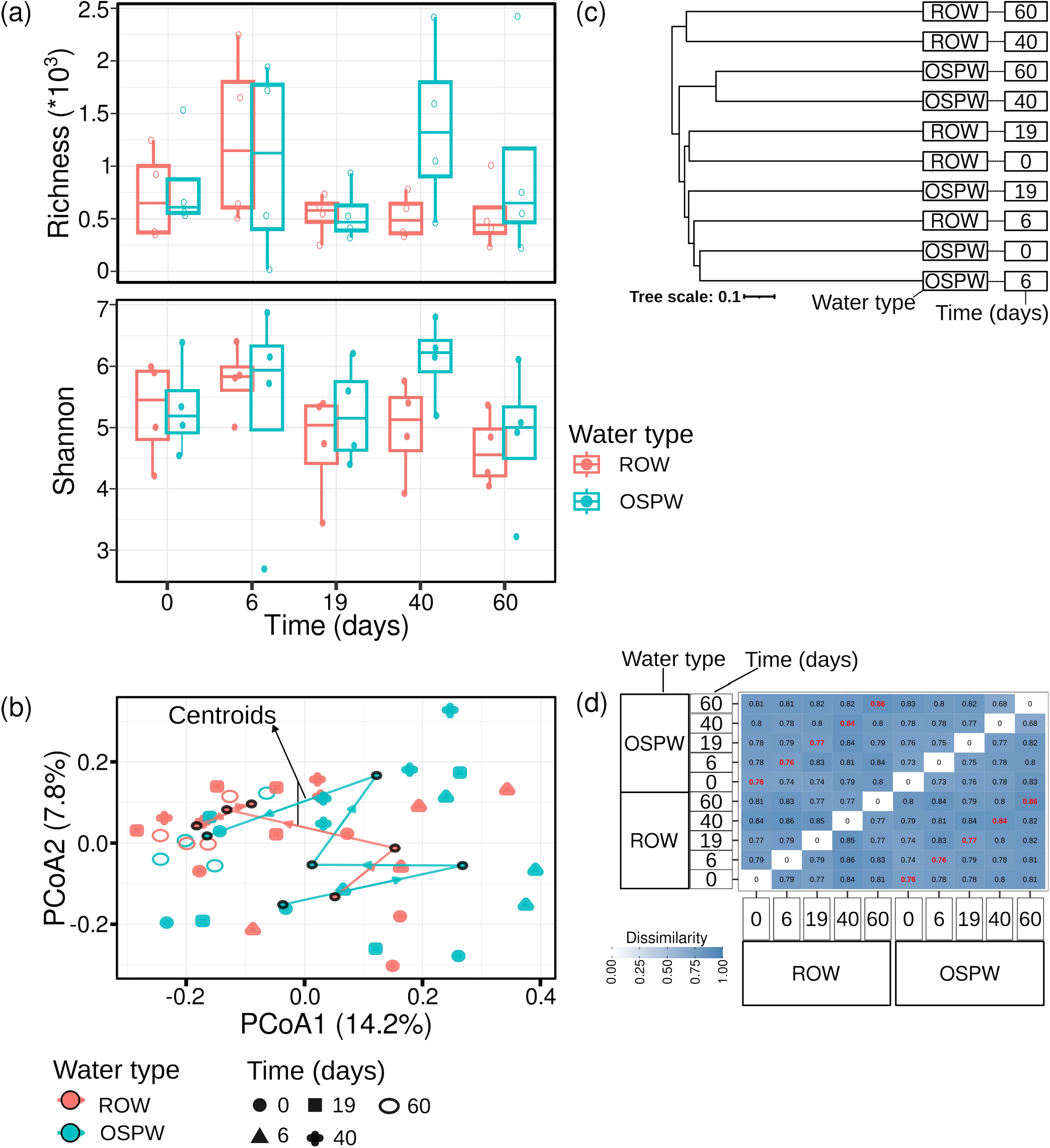
Diversity, UPGMA hierarchical clustering and Bray-Curtis dissimilarity of active microbial communities in root-associated Typha latifolia from mesocosms containing oil sands process-affected water (OSPW) or reverse osmosis water (ROW). (a) Richness and Shannon diversity across water type and time, showing non-significant effects of water type, time and their interaction. (b) Principal Coordinates Analysis (PCoA) based on Bray–Curtis dissimilarity illustrating community succession patterns. Small symbols represent individual replicate samples. Large black-outlined symbols represent group centroids (mean) for each water type and sampling day. Arrows connect centroids chronologically from day 0 to 60. (c) Dendrogram showing sample clustering across time points. (d) Bray-Curtis dissimilarity values between water type at different time points based on the SingleM-derived taxonomic profiles. It ranges from 0.0 (low dissimilarity) to 1.0 (high dissimilarity). Time 0 indicates samples collected before OSPW or fresh ROW treatment period; Time 6-60 represent sampling days after adding OSPW or refilling with ROW.

The pronounced shift at D6 likely reflects a combination of early biological response to the new water chemistry and system-level disturbance associated with draining and refilling. A similar temporal pattern was observed in our DNA-based study of *T. latifolia*, where D40, D60, and D89 samples clustered separately from the earlier time points regardless of water type [14]. In the present metatranscriptomic dataset, Bray-Curtis dissimilarity ranged from 0.76 to 0.86 (Fig. 3b; Table S5), indicating substantial turnover in the active community over time. We therefore interpret the increasing divergence primarily as evidence of strong temporal restructuring, with OSPW contributing an additional selective pressure. The relatively high dissimilarity already present at D0 likely reflects baseline root-to-root heterogeneity, sampling immediately after draining the acclimation water, RNA extraction and technical variation inherent to marker-gene-based active community profiling.

### Plant differentially expressed genes and active detoxification

Because planted OSPW mesocosms in the paired study removed more NAFCs than unplanted controls [14], we examined whether plant roots displayed transcriptional signatures consistent with xenobiotic stress response and detoxification. This is based on the understanding that genes responsive to NAFC belong to broader detoxification systems known to process structurally complex xenobiotics. We therefore analysed plant-derived DEGs to identify *T. latifolia* genes potentially involved in responding to chemically complex OSPW exposure (Fig. 4a; Table S6). Up-regulated genes were identified at D6 (n=3), D19 (n=35), and D40 (n=54), suggesting that the host plant mounted an active molecular response rather than functioning solely as a passive substrate for microbes. These DEGs were mainly classified as oxidoreductases, transferases, carbohydrate transport, plant defense, and transporters (Fig. 4b; Table S7). At D19 and D40, dominant categories included carbohydrate metabolism, oxidoreductases, lipid metabolism and transport, plant defense, transcription, transferases, amino acid metabolism and transport, peptidases and inhibitors, and photosynthesis. As noted above, these responses likely reflect the broader dissolved mixture in OSPW rather than NAFCs alone.

**Fig. 4.**
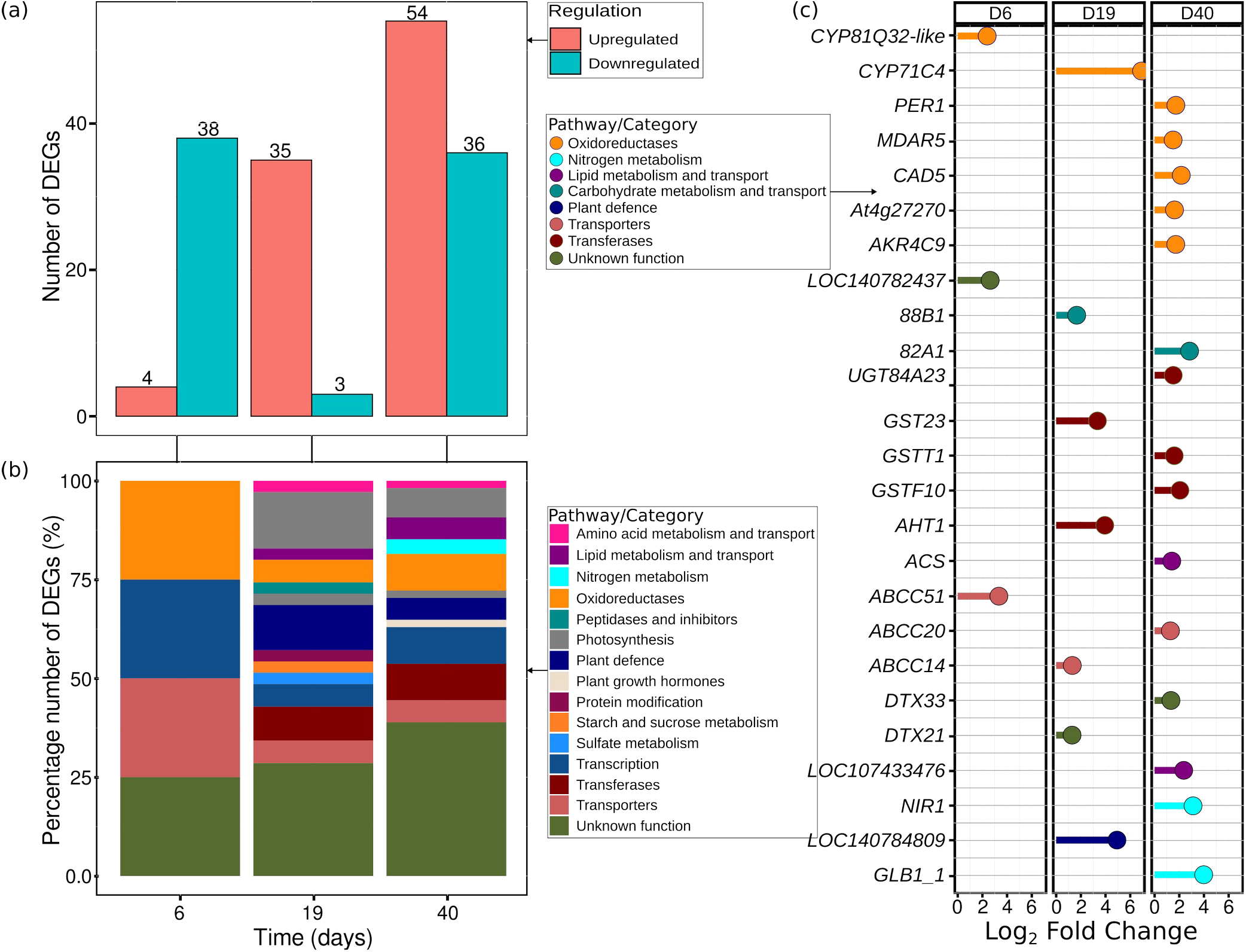
Differentially expressed genes (DEGs) from plant roots. (a) Number of DEGs identified through differential expression analysis in roots grown in mesocosms mimicking constructed wetland systems. (b) Functional categories of the DEGs. (c) Selected up-regulated genes identified in the analysis. Each time (0, 6, 19, 40, and 60) was compared between systems containing oil sands process-affected water (OSPW) and reverse osmosis water (ROW). No DEGs were found at day 60. Time 0 is before addition of OSPW; 6-60 represent sampling days after adding OSPW or refilling with ROW.

We identified 3, 7, and 15 up-regulated plant genesat D6, D19, and D40, respectively, that could plausibly contribute to NAFC-related or broader xenobiotic transformation processes, most of them oxidoreductases (Fig. 4c, Table S7). The responsive DEGs identified in *T. latifolia* were similar to transcripts reported previously in *Arabidopsis thaliana* and *Salix bebbiana* exposed to contaminant stress [13,52]. Plant xenobiotics metabolism is often divided into three different phases: transformation, conjugation, and compartmentalization/exudation [53]. The oxidoreductases identified here are consistent with a phase I-style detoxification response, although their direct substrates cannot be assigned from transcript data alone.

Cytochrome p450s (CYP450s) were identified at the first two time points (D6 and D19), where CYP81Q32-like and CYP71C4 were up-regulated, respectively (Fig. 4c). CYP450s catalyze the oxygenation of diverse xenobiotics in eukaryotes and are often thought to catalyze the first step in xenobiotic metabolism which correlates with our findings [54]. CYP71C4 catalyzes the selective oxygenation or epoxidation of indole derivatives to indoxyl (Phase I detoxification), a precursor of indole alkaloids and other compounds important for plant growth and defense [55,56]. It can also result in indole degradation or unproductive indigo formation [57]. At D40, other oxidoreductases are up-regulated including, peroxidase (PER1), monodehydroascorbate reductase (MDAR5), cinnamyl alcohol dehydrogenase (*CAD5*), NAD(P)H dehydrogenase (quinone; *At4g27270*) and NADPH-dependent aldo-keto reductase (*AKR4C9*). *MDAR5* catalyzes the conversion of monodehydroascorbate to ascorbate while oxidizing NADH, and is known to mediate detoxification of environmental pollutants such as 2,4,6-trinitrotoluene and 1-chloro-2,4-dinitrobenzene [58]. Although alcohol dehydrogenases (*ADHs*) have been proposed to participate in degrading naphthenic acids and polycyclic aromatic compounds in microbes [59,60], this function has not been documented in plants; however, the plant enzyme *CAD5*, which reduces aldehydes to alcohols, may possess similar catalytic potential [61,62]. NAD(P)H-dependent oxidoreductases catalyze a wide variety of redox reactions with many different substrates [63]. *AKR4C9* also acts on a broad range of substrates, including reducing ketosteroids, aromatic aldehydes, ketones, other aliphatic aldehydes, and oxidizes hydroxysteroids [64].

The phase II metabolic encoding genes, uridine diphosphate (UDP) glycosyltransferases (UGTs), were identified at each time point. At D6, 19, and 40 we found that crocetin glucosyltransferase (*LOC140782437*), UDP-glycosyltransferase 88B1-like (*UGT88B1*), and UDP-glycosyltransferase 82A1 (*UGT82A1*) and gallate 1-beta-glucosyltransferase (*UGT84A23*) were up-regulated, respectively (Fig. 4c). The *UGT88B1* gene encodes a protein essential for phase II detoxification, glycosylating lipophilic endogenous and xenobiotic compounds [65]. Further, *UGT84A23* catalyzes the formation of 1-O-β-D-glucose esters from hydroxybenzoic acids and cinnamic acid derivatives [66]. The second most abundant phase II detoxification gene family was glutathione-S-transferases (GSTs) where *GST23* was upregulated at D19 and *GSTT1* and *GSTF10* were upregulated at D40 (Fig. 4c). Both UGTs and GSTs are common phase II detoxification enzymes that link sugars or glutathione to electrophilic functional groups on xenobiotics to increase their water solubility and facilitate transport [76]. GSTs also function in the detoxification of reactive oxygen species, which NAFCs have been reported to produce in plant and animal systems [77,78]. Other up-regulated transferases were identified at D19, agmatine hydroxycinnamoyl transferase 1-like (*AHT1*) and D40, acetyl-coenzyme A synthetase (*ACS*). *AHT1* catalyzes the first step in antifungal hydroxycinnamoyl agmatine biosynthesis by transferring acyl groups from p-coumaroyl-CoA and feruloyl-CoA to acyl acceptors [79,80]. Further, *ACS* has been reported in microbes as part of the degradation pathway for cyclohexene carboxylic acids and may also be involved in the beta-oxidation of some NAFCs [3].

Finally, the last group of genes up-regulated by OSPW were two transporter gene families, the ABC (ATP-binding cassette) transporter type C and the DETOXIFICATION or multidrug and toxic compound efflux carriers (*DTX*/*MATE*). At D6, 19, and 40 the ABCCs 51, 14, and 20 were up-regulated, respectively (Fig. 4c). ABCCs are involved in detoxification of organics and heavy metals, catabolite and phytohormone transport, and ion channel regulation [67]. The *DTX* transporters were less abundant but both *DTX21* and *DTX33* were up-regulated at D19 and D40, respectively (Fig. 4c). Similar to ABC transporters, *DTXs* have been shown to transport organic and inorganic compounds and although *DTX21* does not have a known function, *DTX33* has been shown to transport chloride ions in *Arabidopsis* and this *DTX* may not be involved with NAFC transport [68]. Transporter activity is notable, potentially facilitating NAFC uptake [69]. Other notable DEGs include, 14 kDa proline-rich protein DC2.15-like (*LOC107433476*), ferredoxin-nitrite reductase (*NIR1*), anaerobic nitrite reductase (*GLB1_1*). The proline-rich protein (*LOC107433476*) plays a role in maintaining the strong drought resistance of *Stipa purpurea* population [70]. *NIR1* and *GLB1_1* are involved in nitrate reduction and are noteworthy because nitrate-reducing processes have been implicated in anaerobic NAFC degradation in cooperation with methanogens and sulfate- or iron-reducing microbes [4].

Overall, the substantial up-regulation of detoxification-related genes in *T. latifolia*’s roots supports an active plant response to OSPW exposure. These responses may contribute to the transformation, conjugation, or compartmentalization of organic constituents and could alter the chemical environment experienced by root-associated microbes. However, the present data do not allow us to assign those responses specifically to NAFCs or to conclude that the plant directly prepares compounds for subsequent microbial degradation.

### Microbial differentially expressed genes and their taxonomic origins

The gene expression patterns observed in the microbial community are consistent with the functional differences in NAFC removal between planted and unplanted systems in our previous study **[14]**. In the mesocosms containing plants, NAFC concentrations significantly decreased compared to the unplanted controls. This enhanced removal provides context for interpreting the microbial DEGs, particularly those associated with oxidation–reduction processes and hydrocarbon transformation.

With this in mind, the comparison between OSPW and ROW led to identification of 795 differentially expressed genes (DEGs), with 89% being up-regulated (Fig. 5a; Table S8). The number of DEGs peaked at D40 (354) before sharply declining to D60 (22), coinciding with reduced NAFC concentrations in Balaberda *et al*. [14]. No DEGs were detected at D0, before refilling with distinct water types (ROW or OSPW), which was expected as all mesocosms had been filled with ROW up to that point. Most up-regulated genes were from *Pseudomonadota* (79.3% at D6, declining to 10% at D60), with the *Burkholderiales* and *Rhizobiales* orders as predominant contributors (Fig. 5b & 5c; Table S9–S10). Down-regulated genes were few and taxonomically diverse even among low-abundance orders. Enriched functional categories included translation, ribosomal structure, and stress-related pathways (Fig. S1; Table S11), suggesting protective responses to NAFC toxicity [71].

**Fig. 5.**
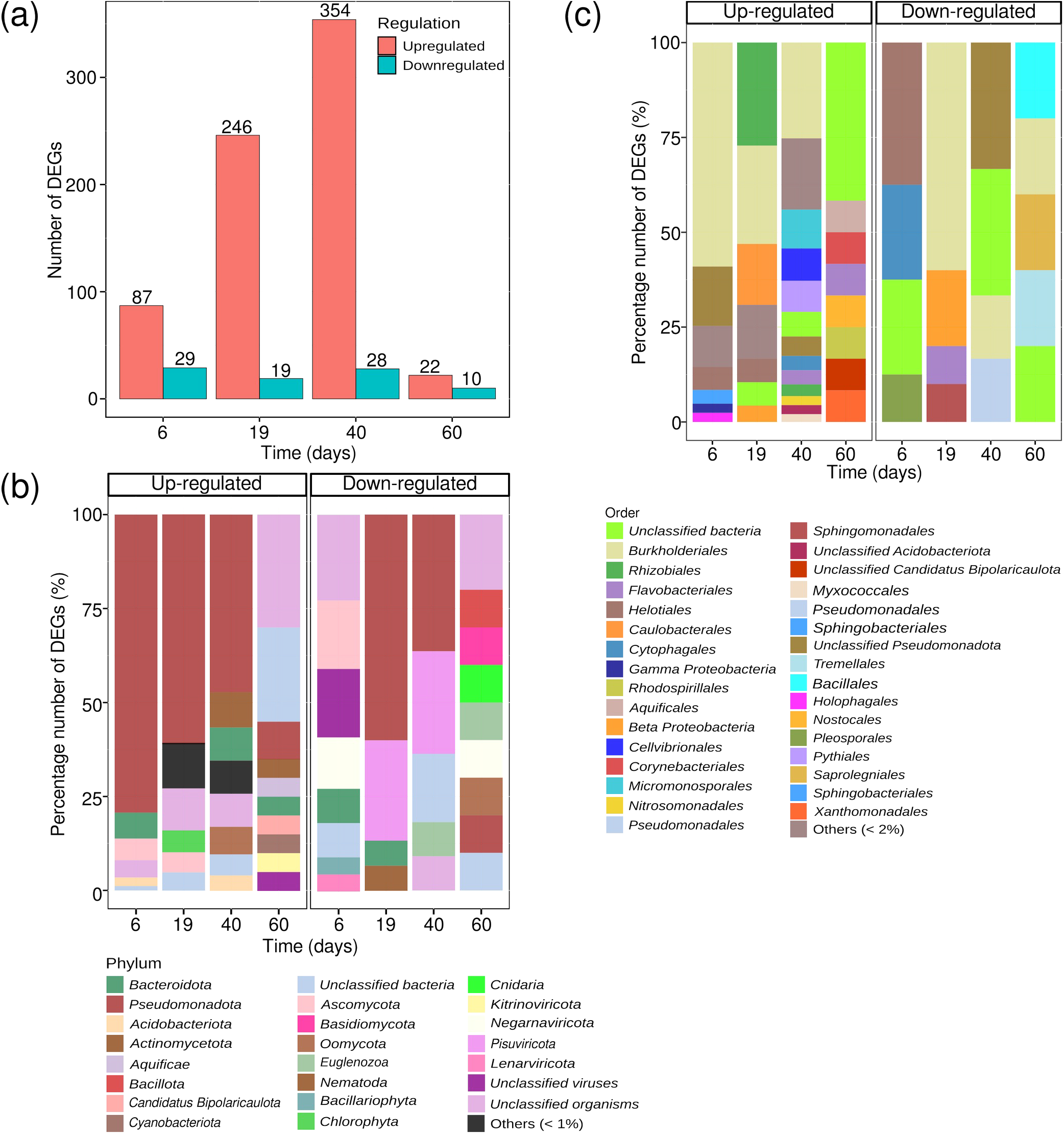
Differentially expressed microbial genes (DEGs) in T. latifolia roots and their taxonomic origins. (a) Number of DEGs at each time point. Taxonomic origins of DEGs at the (b) phylum and (c) order levels. The same time points (0, 6, 19, 40, and 60) were compared between oil sands process-affected water (OSPW) and reverse osmosis water (ROW) treatments. There were no identified DEGs at time 0 before adding OSPW; 6-60 represent sampling days after adding OSPW or refilling with ROW.

Focusing on NAFC-relevant metabolism, 42 up-regulated genes were associated with xenobiotic breakdown across D6, D19, and D40 (Fig. 6; Table S12), including genes involved in oxidoreduction, lipid metabolism, and nutrient (N, P, S) cycling [4,71]. Most of these up-regulated genes were oxidoreductases or transport/metabolism-related enzymes. However, the 18 previously identified NAFC-degradation genes were not found among the up-regulated genes, but we identified specific oxidoreductase genes as reported by Reis *et al. [4]*. These oxidoreductases included NADH-ubiquinone/quinone oxidoreductases (*ND1/4/5, NDUFS2/5 and nuoFGLN*) for xenobiotic detoxification and free radical reduction [60,72]; *cpo* (non-heme chloroperoxidase) for cleaving polymer chains [73] *exaA* (alcohol dehydrogenase)—involved in chloroalkane/chloroalkene degradation [74] *iorB* (isoquinoline 1-oxidoreductase) for targeting nitrogen-containing heterocycles [75] *moxA* (manganese oxidase) for oxidising a variety of phenolic model substrates [76,77]; *prmC* (propane 2-monooxygenase) for degrading propane, acetone, and other volatiles [78,79]; *qor* (NAD(P)H dehydrogenase [quinone]) for reducing and detoxifying quinone and various organics [60,72]; *sucA* (2-oxoglutarate dehydrogenase) for catalysing the oxidative decarboxylation of 2-oxo acids to the corresponding acyl-CoA [80]; *xoxF* (lanthanide-dependent methanol dehydrogenase) for oxidizing alcohols and aldehydes [81]; and *yghU* (GSH-dependent oxidoreductase) for degrading xenobiotics, including lignin, atrazine, and dichloromethane [82]. Additionally, we identified some DEGs as desaturases (*ADS2, desA* and *desC*), which catalyze C–C bond dehydrogenation [83], along with genes involved in lipid metabolism and nitrogen/phosphorus cycling. The genes found here were assigned to *Rhizobiales, Burkholderiales* (*Pseudomonadota*) and *Helotiales* (*Ascomycota*). Together, these results point to a broad oxidative and catabolic response to OSPW rather than a transcriptomic signature that can be assigned uniquely to NAFC degradation.

**Fig. 6.**
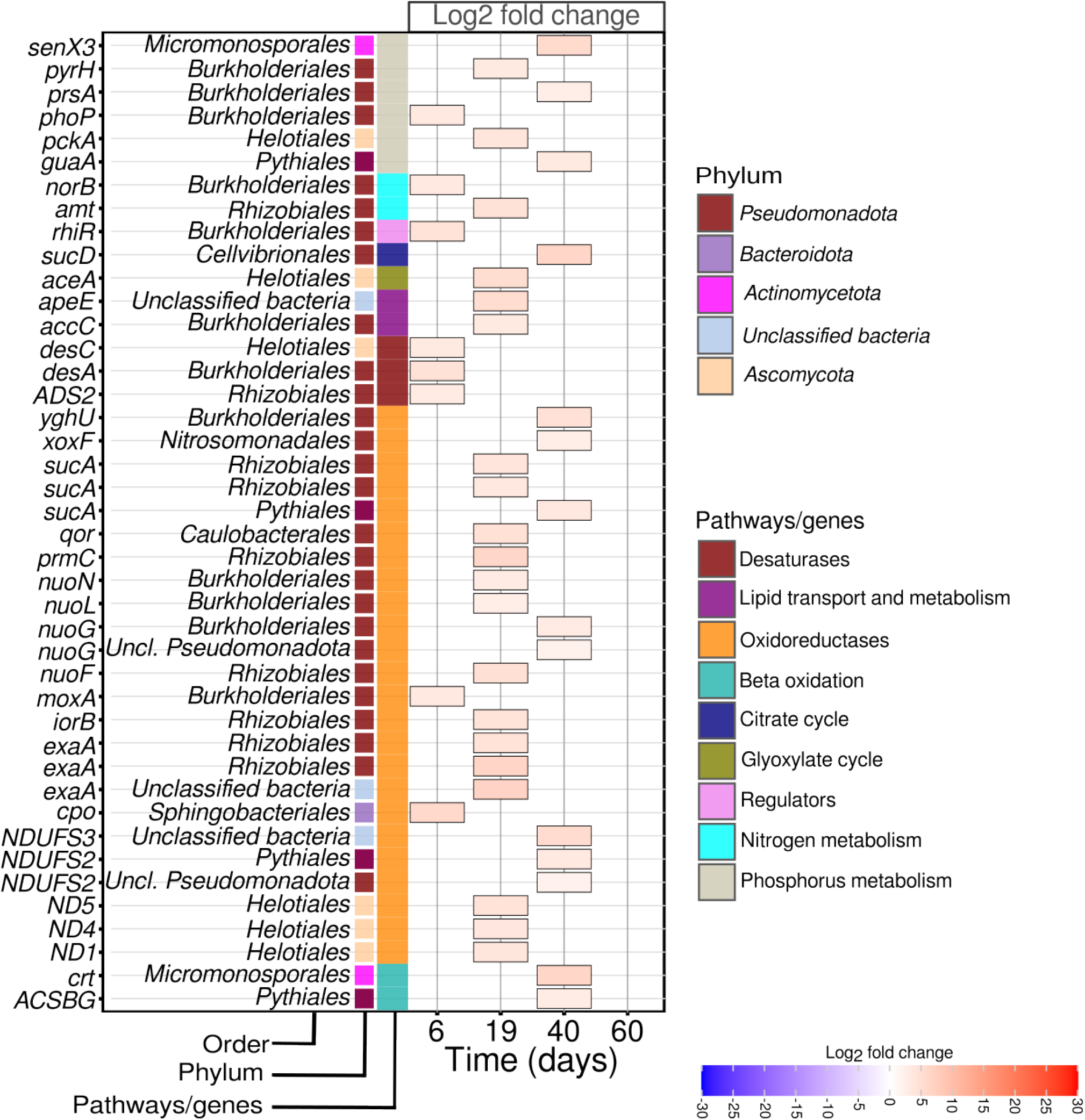
Differentially expressed genes (DEGs) with plausible relevance to naphthenic acid fraction compound transformation or to the metabolism of structurally related hydrocarbons. Comparisons were performed at matched sampling times between oil sands process-affected water (OSPW) and reverse osmosis water (ROW) treatments. There were no identified DEGs at time 0 before adding OSPW; 6-60 represent sampling days after adding OSPW or refilling with ROW.

### Previously reported NAFC-associated genes in the present metatranscriptome

In our microbial differential analysis, the 18 genes previously linked to NAFC degradation [3] were not detected among the DEGs. This is likely because our comparison focused on OSPW versus ROW root samples, rather than the conventional planted versus unplanted rhizosphere mesocosms used in prior studies. Nevertheless, we identified 2,488 sequences across all time points corresponding to previously characterized NAFC-degradation genes, confirming their active expression (Fig. S2; Table S13A). Gene expression peaked at D6 in both OSPW and ROW (sampled after refilling with OSPW or ROW) before declining by D60. The slight peak may be rapid transcriptional response to a slight environmental perturbation. These genes are mostly involved in beta-oxidation pathways. This pattern is consistent with an early response to more bioavailable organic substrates, followed by reduced transcriptional activity as the labile compounds were depleted [84]. Expression was slightly higher after adding OSPW, indicating stimulation by OSPW exposure. Majority of expressed genes were taxonomically assigned to *Pseudomonadota*, which increased after adding or refilling at D6 and declined thereafter (Fig. S3a; Table S13B). This phylum was followed by the pseudofungal phylum *Oomycota*. At the order level, *Burkholderiales* dominated, followed by *Rhizobiales* and *Caulobacterales* (Fig. S4a; Table S13C), consistent with their known roles in NAFCs and organic hydrocarbon degradation [4].

### Abundance and taxonomic origins of hydrocarbon-degradation genes

To explore more NAFC-relevant pathways, we profiled the expression of 37 marker genes from the CANT-HYD database [30], leveraging structural similarities between hydrocarbons and NAFCs. These genes encode enzymes involved in anaerobic and aerobic degradation pathways of aliphatic and aromatic hydrocarbons. Of 37 marker genes, 25 transcripts were recovered, covering 132 sequences across all time points. Transcript abundance generally declined from D6 to D60, except for increases in *prmA* and *tmoA* in ROW (Fig. S5; Table S14A), which encode propane and toluene/benzene monooxygenases essential for oxidation of propane [85] and toluene [86], respectively.

These genes were most abundant at D0 before refilling with OSPW, with other hydrocarbon-degradation genes dominating later, indicating a temporal shift in the hydrocarbon-related transcript pool over time. The same pattern was also observed in hydrocarbon aerobic degradation genes using HADEG (Fig. S6a-c; Table S15A-C), a manually curated database containing sequences of experimentally validated proteins and genes [31], which shares 27 genes with CANT-HYD database. CANT-HYD genes were mainly assigned to *Pseudomonadota, Actinomycetota*, and *Ascomycota* (Fig. S3b; Table S14B), with *Rhizobiales* and *Burkholderiales* as the largest contributors at the order level (Fig. S4b; Table S14C). Similarly, HADEG-derived genes followed similar taxonomic patterns (Fig. S3c; Table S15D), with varying contributions from *Burkholderiales, Rhizobiales*, and *Caulobacterales* (Fig. S4c; Table S15E). These orders are among the key taxa reported to possess the genes for NAFC degradation [4].

### Abundance and taxonomic origins of nutrient cycling genes

We profiled genes that could be linked to anaerobic NAFC degradation, including those linked to nitrogen, sulfur, phosphorus, and methanogenic pathways [87,88]. In general, there were no consistent temporal patterns in genes involved in N, S, or P metabolism across systems (Fig. S7–S10; Table S16-S19A-B). However, OSPW demonstrated a rise in sulfur-related genes at D6 (Fig. S8; Table S17A–B). These genes were primarily assigned to *Pseudomonadota, Bacteroidota, Ascomycota*, and *Oomycota* (Fig. S11A-D; Table S16C–S19C), with *Burkholderiales* and *Rhizobiales* consistently dominating among orders (Fig. S12; Table S16D–S19D). Both groups are well known for nitrate reduction, denitrification, sulfur cycling, and phosphorus mineralization [89–92], supporting their potential role in transforming heteroatom-containing organic constituents in OSPW. Methanogenesis was unlikely, as key *mcrABCDG* genes were absent (Fig. S10; Table S19); instead, expressed genes were linked to methylotrophy rather than methane production [93], with contributions from *Rhizobiales, Burkholderiales*, and other methylotrophic taxa.

## Conclusion

Our findings show coordinated plant and microbial transcriptional responses in *Typha latifolia-*planted mesocosms previously shown to remove NAFCs from OSPW. The active microbiome was initially dominated by *Pseudomonadota*, especially Burkholderiales, and shifted over time alongside broad changes in functional gene expression. In pa*rallel, T. latifolia* roots up-regulated genes associated with oxidoreduction, transport, and conjugation, consistent with xenobiotic stress responses. Overall, these results help explain the stronger NAFC attenuation previously observed in planted mesocosms, but they do not by themselves show direct NAFC degradation by particular taxa or demonstrate plant-microbe synergy. Because OSPW is chemically complex, the transcriptional patterns observed here likely reflect combined responses to NAFCs and other dissolved constituents. Future work should pair transcriptomics with time-resolved chemistry, isolate-based validation, and plant-only or microbe-only controls to directly test the mechanisms suggested here.

## Availability of data and materials

The metatranscriptomic fasta files were deposited under the NCBI BioProject PRJNA1358281 (see Table S20). For reproducibility, reusability, and transparency, we have also made available the co-assembled library (doi:10.6084/m9.figshare.30667601). All scripts used after data analysis and figure generation are publicly available on GitHub (https://github.com/Julius-Nweze/Exp3_differential_expression_analysis/tree/v1.0) or Zenodo (https://doi.org/10.5281/zenodo.17553524)

## Acknowledgements

We acknowledge funding support from a Genome Canada Large-Scale Applied Research Project (LSARP) grant (#18207), in partnership with Genome Alberta and Genome Québec. We also thank Imperial Oil Resources Limited for providing OSPW, Kaitlyn Carey and Kaela Walton-Sather for their assistance in the greenhouse and Marie-Josée Morency for her assistance with library preparation.

## Contributions

Degenhardt, Muench, Yergeau, and Martineau conceptualized the experiments and overall approach. Bergeron supervised and conducted laboratory analyses. Yergeau, Nweze, Morvan and Samad designed the data analyses, and Nweze performed the bioinformatics and statistical analyses, with contributions from Tremblay (bioinformatics), and Muench and Symonds (analysis and interpretation) for the plant transcriptome data analysis. Nweze wrote the manuscript with substantial input from Yergeau, Morvan, Samad and Symonds. All authors read and approved the final version of the manuscript.

## References

1. Li C, Fu L, Stafford J et al. The toxicity of oil sands process-affected water (OSPW): A critical review. Sci Total Environ 2017;601–602:1785–802. 10.1016/j.scitotenv.2017.06.024.

2. Vander Meulen IJ, Schock DM, Parrott JL et al. Characterization of naphthenic acid fraction compounds in water from Athabasca oil sands wetlands by Orbitrap high-resolution mass spectrometry. Sci Total Environ 2021;780:146342. 10.1016/j.scitotenv.2021.146342.

3. Quinlan PJ, Tam KC. Water treatment technologies for the remediation of naphthenic acids in oil sands process-affected water. Chem Eng J 2015;279:696–714. 10.1016/j.cej.2015.05.062.

4. Reis PCJ, Correa-Garcia S, Tremblay J et al. Microbial degradation of naphthenic acids using constructed wetland treatment systems: metabolic and genomic insights for improved bioremediation of process-affected water. FEMS Microbiol Ecol 2023;99(12):fiad153. 10.1093/femsec/fiad153.

5. Das N, Chandran P. Microbial degradation of petroleum hydrocarbon contaminants: An overview. Biotechnol Res Int 2011;2011(1):941810. 10.4061/2011/941810.

6. Varjani SJ. Microbial degradation of petroleum hydrocarbons. Bioresour Technol 2017;223:277–86. 10.1016/j.biortech.2016.10.037.

7. Bhandari S, Poudel DK, Marahatha R et al. Microbial Enzymes used in bioremediation. J Chem 2021;2021(1):8849512. 10.1155/2021/8849512.

8. Yergeau E, Lawrence JR, Sanschagrin S et al. Next-generation sequencing of microbial communities in the Athabasca River and its tributaries in relation to oil sands mining activities. Appl Environ Microbiol 2012;78(21):7626–37. 10.1128/AEM.02036-12.

9. Mitter EK, Freitas JR de, Germida JJ. Bacterial root microbiome of plants growing in oil sands reclamation covers. Front Microbiol 2017;8:849. 10.3389/fmicb.2017.00849.

10. Alberts ME, Wong J, Hindle R et al. Detection of naphthenic acid uptake into root and shoot tissues indicates a direct role for plants in the remediation of oil sands process-affected water. Sci Total Environ 2021;795:148857. 10.1016/j.scitotenv.2021.148857.

11. Santoyo G. How plants recruit their microbiome? New insights into beneficial interactions. J Adv Res 2021;40:45–58. 10.1016/j.jare.2021.11.020.

12. Rodriguez-Hernandez MC, García De la-Cruz RF, Leyva E et al. Typha latifolia as potential phytoremediator of 2,4-dichlorophenol: Analysis of tolerance, uptake and possible transformation processes. Chemosphere 2017;173:190–8. 10.1016/j.chemosphere.2016.12.043.

13. Widdup EE, Chatfield-Reed K, Henry D et al. Identification of detoxification pathways in plants that are regulated in response to treatment with organic compounds isolated from oil sands process-affected water. Chemosphere 2015;139:47–53. 10.1016/j.chemosphere.2015.05.048.

14. Balaberda A lynne, Nweze JE, Meulen IJV et al. Role of Typha latifolia, Juncus balticus, and root-associated microbial communities in naphthenic acid removal from oil sands process-affected water. Wetlands 2026:4–7.

15. Parada AE, Needham DM, Fuhrman JA. Every base matters: assessing small subunit rRNA primers for marine microbiomes with mock communities, time series and global field samples. Environ Microbiol 2016;18(5):1403–14. 10.1111/1462-2920.13023.

16. Quince C, Lanzen A, Davenport RJ et al. Removing noise from pyrosequenced amplicons. BMC Bioinformatics 2011;12:38. 10.1186/1471-2105-12-38.

17. Woodcroft BJ, Aroney STN, Zhao R et al. Comprehensive taxonomic identification of microbial species in metagenomic data using SingleM and Sandpiper. Nat Biotechnol published online 16 July 2025. 10.1038/s41587-025-02738-1.

18. Loewenstein Y, Portugaly E, Fromer M et al. Efficient algorithms for accurate hierarchical clustering of huge datasets: tackling the entire protein space. Bioinformatics 2008;24(13):i41–9. 10.1093/bioinformatics/btn174.

19. Tremblay J, Yergeau E, Fortin N et al. Chemical dispersants enhance the activity of oil- and gas condensate-degrading marine bacteria. Int Soc Microb Ecol J 2017;11(12):2793–808. 10.1038/ismej.2017.129.

20. Bourgey M, Dali R, Eveleigh R et al. GenPipes: an open-source framework for distributed and scalable genomic analyses. GigaScience 2019;8(6):giz037. 10.1093/gigascience/giz037.

21. Bolger AM, Lohse M, Usadel B. Trimmomatic: a flexible trimmer for Illumina sequence data. Bioinformatics 2014;30(15):2114–20. 10.1093/bioinformatics/btu170.

22. Li D, Liu CM, Luo R et al. MEGAHIT: an ultra-fast single-node solution for large and complex metagenomics assembly via succinct de Bruijn graph. Bioinformatics 2015;31(10):1674–6. 10.1093/bioinformatics/btv033.

23. Li H. Aligning sequence reads, clone sequences and assembly contigs with BWA-MEM, 1303.3997. Preprint, arXiv, 26 May 2013. 10.48550/arXiv.1303.3997.

24. Dobin A, Davis CA, Schlesinger F et al. STAR: ultrafast universal RNA-seq aligner. Bioinforma Oxf Engl 2013;29(1):15–21. 10.1093/bioinformatics/bts635.

25. Love MI, Huber W, Anders S. Moderated estimation of fold change and dispersion for RNA-seq data with DESeq2. Genome Biol 2014;15(12):550. 10.1186/s13059-014-0550-8.

26. Benjamini Y, Hochberg Y. Controlling the false discovery rate: a practical and powerful approach to multiple testing. J R Stat Soc Ser B Methodol 1995;57(1):289–300. 10.1111/j.2517-6161.1995.tb02031.x.

27. Hyatt D, Chen GL, LoCascio PF et al. Prodigal: prokaryotic gene recognition and translation initiation site identification. BMC Bioinformatics 2010;11(1):119. 10.1186/1471-2105-11-119.

28. Johnson LS, Eddy SR, Portugaly E. Hidden Markov model speed heuristic and iterative HMM search procedure. BMC Bioinformatics 2010;11(1):431. 10.1186/1471-2105-11-431.

29. Kanehisa M, Goto S. KEGG: kyoto encyclopedia of genes and genomes. Nucleic Acids Res 2000;28(1):27–30. 10.1093/nar/28.1.27.

30. Khot V, Zorz J, Gittins DA et al. CANT-HYD: A curated database of phylogeny-derived hidden markov models for annotation of marker genes involved in hydrocarbon degradation. Front Microbiol 2022;12. 10.3389/fmicb.2021.764058.

31. Rojas-Vargas J, Castelán-Sánchez HG, Pardo-López L. HADEG: A curated hydrocarbon aerobic degradation enzymes and genes database. Comput Biol Chem 2023;107:107966. 10.1016/j.compbiolchem.2023.107966.

32. Qian L, Yu X, Zhou J et al. MCycDB: A curated database for comprehensively profiling methane cycling processes of environmental microbiomes. Mol Ecol Resour 2022;22(5):1803–23. 10.1111/1755-0998.13589.

33. Tu Q, Lin L, Cheng L et al. NCycDB: a curated integrative database for fast and accurate metagenomic profiling of nitrogen cycling genes. Bioinformatics 2019;35(6):1040–8. 10.1093/bioinformatics/bty741.

34. Yu X, Zhou J, Song W et al. SCycDB: A curated functional gene database for metagenomic profiling of sulphur cycling pathways. Mol Ecol Resour 2021;21(3):924–40. 10.1111/1755-0998.13306.

35. Zeng J, Tu Q, Yu X et al. PCycDB: a comprehensive and accurate database for fast analysis of phosphorus cycling genes. Microbiome 2022;10(1):101. 10.1186/s40168-022-01292-1.

36. Ye J, McGinnis S, Madden TL. BLAST: improvements for better sequence analysis. Nucleic Acids Res 2006;34(Web Server issue):W6–9. 10.1093/nar/gkl164.

37. R Core Team. R: A language and environment for statistical computing. R Foundation for Statistical Computing, Vienna, Austria, 2021. https://www.R-project.org/.

38. Wickham H, Wickham H. Programming with Ggplot2: Elegant Graphics for Data Analysis. Springer, 2016. https://ggplot2-book.org/.

39. Letunic I, Bork P. Interactive tree of life (iTOL) v5: an online tool for phylogenetic tree display and annotation. Nucleic Acids Res 2021;49(W1):W293–6. 10.1093/nar/gkab301.

40. Zymela A, Muench DG. Optimized Method for Cultivation and Microbial Bioaugmentation of Typha latifolia (Cattail). J Vis Exp JoVE July 2025;(221). 10.3791/67729.

41. Rodriguez-Hernandez MC, García De la-Cruz RF, Leyva E et al. Typha latifolia as potential phytoremediator of 2,4-dichlorophenol: Analysis of tolerance, uptake and possible transformation processes. Chemosphere 2017;173:190–8. 10.1016/j.chemosphere.2016.12.043.

42. Zapata-Morales AL, Alfaro-De la Torre MaC, Hernández-Morales A et al. Isolation of Cultivable Bacteria Associated with the Root of Typha latifolia in a Constructed Wetland for the Removal of Diclofenac or Naproxen. Water Air Soil Pollut 2020;231(8):423. 10.1007/s11270-020-04781-x.

43. Singleton DR, Adrion AC, Aitken MD. Surfactant-induced bacterial community changes correlated with increased polycyclic aromatic hydrocarbon degradation in contaminated soil. Appl Microbiol Biotechnol 2016;100(23):10165–77. 10.1007/s00253-016-7867-z.

44. Liu T, Lan L, Li Y et al. Radial oxygen loss of Typha latifolia outperforms microbial effects in heavy metal(loid) stabilization. Environ Res 2025;285:122561. 10.1016/j.envres.2025.122561.

45. Pietrangelo L, Bucci A, Maiuro L et al. Unraveling the composition of the root-associated bacterial microbiota of Phragmites australis and Typha latifolia. Front Microbiol 2018;9. 10.3389/fmicb.2018.01650.

46. Chi H, Yang L, Yang W et al. Variation of the bacterial community in the rhizoplane iron plaque of the wetland plant Typha latifolia. Int J Environ Res Public Health 2018;15(12):2610. 10.3390/ijerph15122610.

47. Samad A, Degenhardt D, Seguin A et al. Microbial community structural and functional differentiation in capped thickened oil sands tailings planted with native boreal species. Front Microbiol 2023;14:1168653. 10.3389/fmicb.2023.1168653.

48. Rojas-Rojas FU, Gómez-Vázquez IM, Estrada-de los Santos P et al. The potential of Paraburkholderia species to enhance crop growth. World J Microbiol Biotechnol 2025;41(2):62. 10.1007/s11274-025-04256-3.

49. Seo H, Kim JH, Lee SM et al. The Plant-associated Flavobacterium: A hidden helper for improving plant health. Plant Pathol J 2024;40(3):251–60. 10.5423/PPJ.RW.01.2024.0019.

50. Liu N, Liu Q, Min J et al. Specific bacterial communities in the rhizosphere of low-cadmium and high-zinc wheat (Triticum aestivum L.). Sci Total Environ 2022;838:156484. 10.1016/j.scitotenv.2022.156484.

51. Martínez-Martínez JG, Rosales-Loredo S, Hernández-Morales A et al. Bacterial communities associated with the roots of Typha spp. and Its relationship in phytoremediation processes. Microorganisms 2023;11(6):1587. 10.3390/microorganisms11061587.

52. Armstrong SA, Headley JV, Peru KM et al. Differences in phytotoxicity and dissipation between ionized and nonionized oil sands naphthenic acids in wetland plants. Environ Toxicol Chem 2009;28(10):2167–74. 10.1897/09-059.1.

53. Samad A, Pelletier G, Séguin A et al. Understanding willow transcriptional response in the context of oil sands tailings reclamation. Front Plant Sci 2022;13. 10.3389/fpls.2022.857535.

54. Kaur S, Rani S, Mishra S et al. Mechanistic understanding of xenobiotic detoxification by plants. In: Roy S, Mandal V (eds), Plant-Microbe Interaction under Xenobiotic Exposure. Singapore: Springer Nature, 2025, 243–57. 10.1007/978-981-96-8260-7_8.

55. Behrendorff JBYH. Reductive cytochrome P450 reactions and their potential Role in bioremediation. Front Microbiol 2021;12. 10.3389/fmicb.2021.649273.

56. Heine T, Großmann C, Hofmann S et al. Indigoid dyes by group E monooxygenases: mechanism and biocatalysis. Biol Chem 2019;400(7):939–50. 10.1515/hsz-2019-0109.

57. Sorg A, Luo ZW, Li QB et al. The airborne herbivore-induced plant volatile indole is converted to benzoxazinoid defense compounds in maize plants. New Phytol 2025;246(2):718–28. 10.1111/nph.70004.

58. Tischler D, Kumpf A, Eggerichs D et al. Chapter Thirteen -Styrene monooxygenases, indole monooxygenases and related flavoproteins applied in bioremediation and biocatalysis. In: Chaiyen P, Tamanoi F (eds), The Enzymes, Flavin-Dependent Enzymes: Mechanisms, Structures and Applications. n.p.: Academic Press, 2020, 47.399–425. 10.1016/bs.enz.2020.05.011.

59. Johnston EJ, Rylott EL, Beynon E et al. Monodehydroascorbate reductase mediates TNT toxicity in plants. Science 2015;349(6252):1072–5. 10.1126/science.aab3472.

60. McKew BA, Johnson R, Clothier L et al. Differential protein expression during growth on model and commercial mixtures of naphthenic acids in Pseudomonas fluorescens Pf-5. MicrobiologyOpen 2021;10(4):e1196. 10.1002/mbo3.1196.

61. Naveed M, Iqbal F, Aziz T et al. Exploration of alcohol dehydrogenase EutG from Bacillus tropicus as an eco-friendly approach for the degradation of polycyclic aromatic compounds. Sci Rep 2025;15(1):3466. 10.1038/s41598-025-86624-5.

62. Sibout R, Eudes A, Pollet B et al. Expression pattern of two paralogs encoding cinnamyl alcohol dehydrogenases in Arabidopsis. Isolation and characterization of the corresponding mutants. Plant Physiol 2003;132(2):848–60. 10.1104/pp.103.021048.

63. Kim SJ, Kim MR, Bedgar DL et al. Functional reclassification of the putative cinnamyl alcohol dehydrogenase multigene family in Arabidopsis. Proc Natl Acad Sci U S A 2004;101(6):1455–60. 10.1073/pnas.0307987100.

64. Sellés Vidal L, Kelly CL, Mordaka PM et al. Review of NAD(P)H-dependent oxidoreductases: Properties, engineering and application. Biochim Biophys Acta BBA - Proteins Proteomics 2018;1866(2):327–47. 10.1016/j.bbapap.2017.11.005.

65. Simpson PJ, Tantitadapitak C, Reed AM et al. Characterization of two novel aldo-keto reductases from Arabidopsis: expression patterns, broad substrate specificity, and an open active-site structure suggest a role in toxicant metabolism following stress. J Mol Biol 2009;392(2):465–80. 10.1016/j.jmb.2009.07.023.

66. Asif MZ, Nocilla KA, Ngo L et al. Role of UDP-Glycosyltransferase (ugt) genes in detoxification and glycosylation of 1-hydroxyphenazine (1-HP) in Caenorhabditis elegans. Chem Res Toxicol 2024;37(4):590–9. 10.1021/acs.chemrestox.3c00410.

67. Tanabe K, Hojo Y, Shinya T et al. Molecular evidence for biochemical diversification of phenolamide biosynthesis in rice plants. J Integr Plant Biol 2016;58(11):903–13. 10.1111/jipb.12480.

68. Uragami T, Kiba T, Kojima M et al. The cytokinin efflux transporter ABCC4 participates in Arabidopsis root system development. Plant Physiol 2024;197(1):kiae628. 10.1093/plphys/kiae628.

69. Zhang L, Xu Q, Zhan S et al. A new NAD(H)-dependent meso-2,3-butanediol dehydrogenase from an industrially potential strain Serratia marcescens H30. Appl Microbiol Biotechnol 2014;98(3):1175–84. 10.1007/s00253-013-4959-x.

70. Zhang H, Zhao FG, Tang RJ et al. Two tonoplast MATE proteins function as turgor-regulating chloride channels in Arabidopsis. Proc Natl Acad Sci 2017;114(10):E2036–45. 10.1073/pnas.1616203114.

71. Li X, Yang Y, Yang S et al. Comparative proteomics analyses of intraspecific differences in the response of Stipa purpurea to drought. Plant Divers 2016;38(2):101–17. 10.1016/j.pld.2016.03.002.

72. Lou F, Okoye CO, Gao L et al. Whole-genome sequence analysis reveals phenanthrene and pyrene degradation pathways in newly isolated bacteria Klebsiella michiganensis EF4 and Klebsiella oxytoca ETN19. Microbiol Res 2023;273:127410. 10.1016/j.micres.2023.127410.

73. Pey AL, Megarity CF, Timson DJ. NAD(P)H quinone oxidoreductase (NQO1): an enzyme which needs just enough mobility, in just the right places. Biosci Rep 2019;39(1):BSR20180459. 10.1042/BSR20180459.

74. Rong Z, Ding ZH, Wu YH et al. Degradation of low-density polyethylene by the bacterium Rhodococcus sp. C-2 isolated from seawater. Sci Total Environ 2024;907:167993. 10.1016/j.scitotenv.2023.167993.

75. Elumalai P, Parthipan P, Huang M et al. Enhanced biodegradation of hydrophobic organic pollutants by the bacterial consortium: Impact of enzymes and biosurfactants. Environ Pollut 2021;289:117956. 10.1016/j.envpol.2021.117956.

76. Kotoky R, Ogawa N, Pandey P. The structure-function relationship of bacterial transcriptional regulators as a target for enhanced biodegradation of aromatic hydrocarbons. Microbiol Res 2022;262:127087. 10.1016/j.micres.2022.127087.

77. Sherif M, Waung D, Korbeci B et al. Biochemical studies of the multicopper oxidase (small laccase) from Treptomyces coelicolor using bioactive phytochemicals and site-directed mutagenesis. Microb Biotechnol 2013;6(5):588–97. 10.1111/1751-7915.12068.

78. Romano CA, Zhou M, Song Y et al. Biogenic manganese oxide nanoparticle formation by a multimeric multicopper oxidase Mnx. Nat Commun 2017;8(1):746. 10.1038/s41467-017-00896-8.

79. Furuya T, Nakao T, Kino K. Catalytic function of the mycobacterial binuclear iron monooxygenase in acetone metabolism. FEMS Microbiol Lett 2015;362(19):fnv136. 10.1093/femsle/fnv136.

80. Furuya T, Hirose S, Osanai H et al. Identification of the Monooxygenase gene clusters responsible for the regioselective oxidation of phenol to hydroquinone in Mycobacteria. Appl Environ Microbiol 2011;77(4):1214–20. 10.1128/AEM.02316-10.

81. Heath C, Posner MG, Aass HC et al. The 2-oxoacid dehydrogenase multi-enzyme complex of the archaeon Thermoplasma acidophilum − recombinant expression, assembly and characterization. Fed Eur Biochem Soc 2007;274(20):5406–15. 10.1111/j.1742-4658.2007.06067.x.

82. Wehrmann M, Billard P, Martin-Meriadec A et al. Functional role of lanthanides in enzymatic activity and transcriptional regulation of pyrroloquinoline quinone-dependent alcohol dehydrogenases in Pseudomonas putida KT2440. mBio 2017;8(3):10.1128/mbio.00570-17. https://doi.org/10.1128/mbio.00570-17.

83. Allocati N, Federici L, Masulli M et al. Glutathione transferases in bacteria. Fed Eur Biochem Soc 2009;276(1):58–75. 10.1111/j.1742-4658.2008.06743.x.

84. Cerone M, Smith TK. Desaturases: Structural and mechanistic insights into the biosynthesis of unsaturated fatty acids. IUBMB Life 2022;74(11):1036–51. 10.1002/iub.2671.

85. Herman DC, Fedorak PM, MacKinnon MD et al. Biodegradation of naphthenic acids by microbial populations indigenous to oil sands tailings. Can J Microbiol 1994;40(6):467–77. 10.1139/m94-076.

86. Kotani T, Yamamoto T, Yurimoto H et al. Propane monooxygenase and NAD+-dependent secondary alcohol dehydrogenase in propane metabolism by Gordonia sp. strainTY-5. J Bacteriol 2003;185(24):7120–8. 10.1128/jb.185.24.7120-7128.2003.

87. Whited GM, Gibson DT. Toluene-4-monooxygenase, a three-component enzyme system that catalyzes the oxidation of toluene to p-cresol in Pseudomonas mendocina KR1. J Bacteriol 1991;173(9):3010–6. 10.1128/jb.173.9.3010-3016.1991.

88. Clothier LN, Gieg LM. Anaerobic biodegradation of surrogate naphthenic acids. Water Res 2016;90:156–66. 10.1016/j.watres.2015.12.019.

89. Kinley CM, Gaspari DP, McQueen AD et al. Effects of environmental conditions on aerobic degradation of a commercial naphthenic acid. Chemosphere 2016;161:491–500. 10.1016/j.chemosphere.2016.07.050.

90. Łochowska A, Iwanicka-Nowicka R, Zielak A et al. Regulation of sulfur assimilation pathways in Burkholderia cenocepacia through control of genes by the ssuR transcription factor▿. J Bacteriol 2011;193(8):1843–53. 10.1128/JB.00483-10.

91. Ragot SA, Huguenin-Elie O, Kertesz MA et al. Total and active microbial communities and phoD as affected by phosphate depletion and pH in soil. Plant Soil 2016;408(1):15–30. 10.1007/s11104-016-2902-5.

92. Gan X, Hu H, Fu Q et al. Nitrate reduction coupling with As(III) oxidation in neutral As-contaminated paddy soil preserves nitrogen, reduces N_2_O emissions and alleviates As toxicity. Sci Total Environ 2024;912:169360. 10.1016/j.scitotenv.2023.169360.

93. Ramírez-Amador F, Paul S, Kumar A et al. Structure of the ATP-driven methyl-coenzyme M reductase activation complex. Nature 2025;642(8068):814–21. 10.1038/s41586-025-08890-7.

