## Supplementary Figures for "Coordinated plant and microbial transcriptional responses to oil-sands process-affected water"

### Short title: Insights from *Typha latifolia* root metatranscriptomes in constructed wetland mesocosms

**Nweze J.E[a], Morvan S.[a], Samad A.[a], Bergeron M-J.[b], Degenhardt D.[c], Tremblay J.[d], Symonds K.[e], Muench D.G.[e], Martineau C.[b] and Yergeau E.[a]***

[a]Institut National de Recherche Scientifique, Centre Armand Frappier Santé et Biotechnologie, Laval, QC, Canada

[b]Natural Resources Canada, Canadian Forest Service, Laurentian Forestry Centre, Québec, QC, Canada

[c]Natural Resources Canada, Canadian Forest Service, Northern Forestry Centre, Edmonton, AB, Canada

[d]Astek Canada, Montréal, QC, Canada

[e]University of Calgary, Department of Biological Sciences, Calgary, AB, Canada

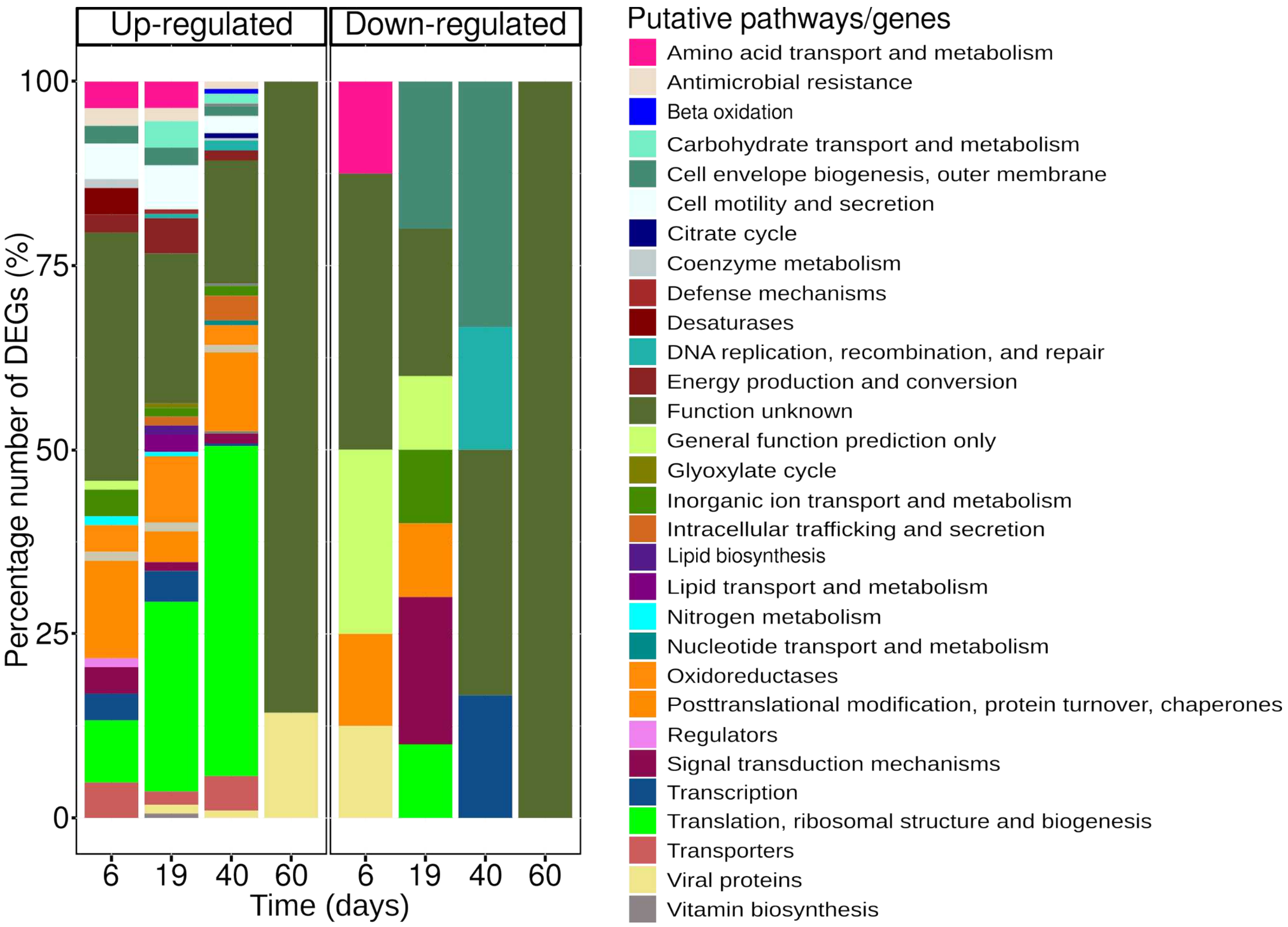


*Fig. S1 Pathways/categories of the differentially expressed genes (DEGs) from plant roots. The figure represents the metabolic pathway or gene categories of DEGs identified through differential expression analysis in T. latifolia grown in mesocosms mimicking constructed wetland systems. The same time points (0, 6, 19, 40, and 60) were compared between oil sands process-affected water*

*(OSPW) and reverse osmosis water (ROW). Time 0 is before addition of OSPW; 6-60 represent sampling days after adding OSPW or refilling with ROW.*

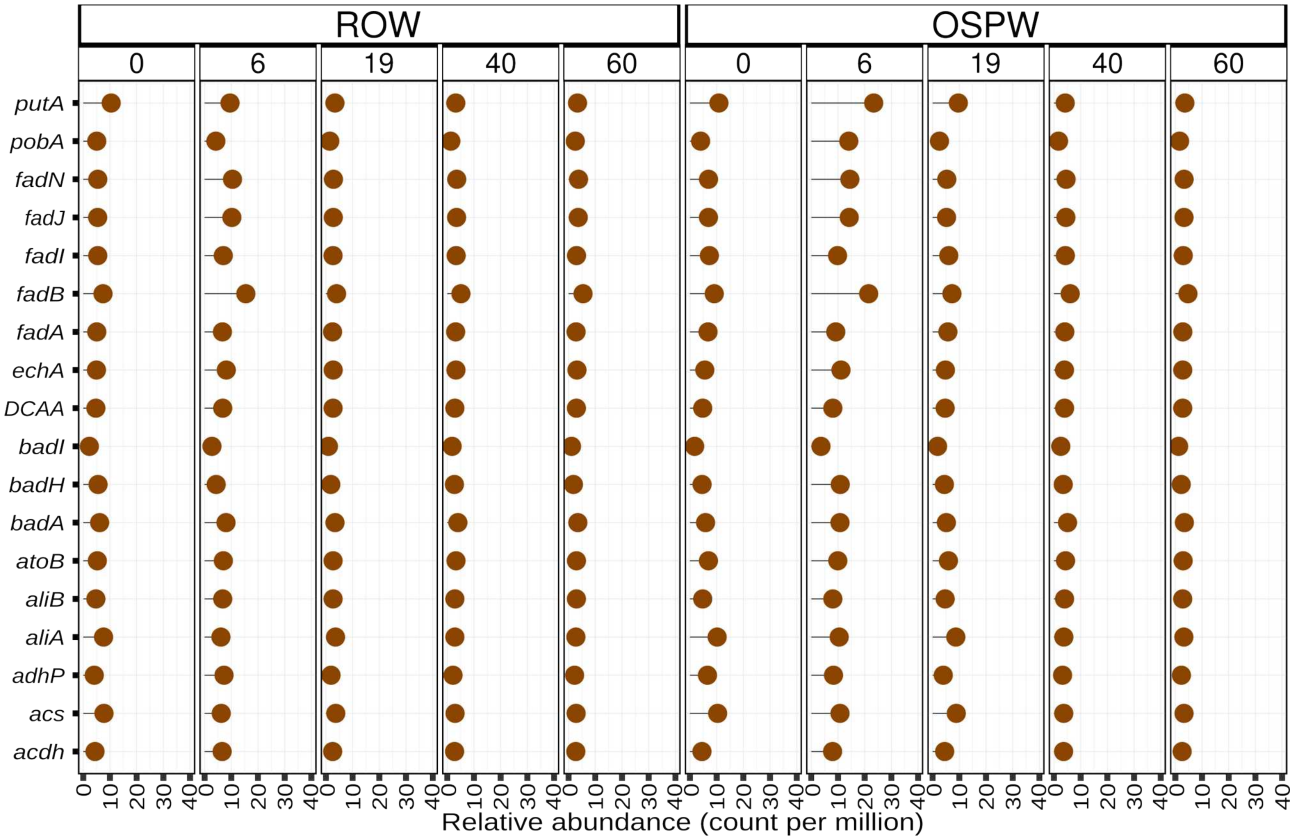


***Fig. S2. Abundance of previously identified naphthenic-acid-degrading genes (NA-genes).*** *The figure represents the average abundance of NA-genes calculated in count per million (CPM). The data came from the root of T. latifolia grown in mesocosms mimicking constructed wetland systems containing either reverse osmosis water (ROW) or oil sands process-affected water (OSPW) at different time points (0, 6, 19, 40, and 60). Time 0 is before addition of OSPW; 6-60 represent sampling days after adding OSPW or refilling with ROW.*

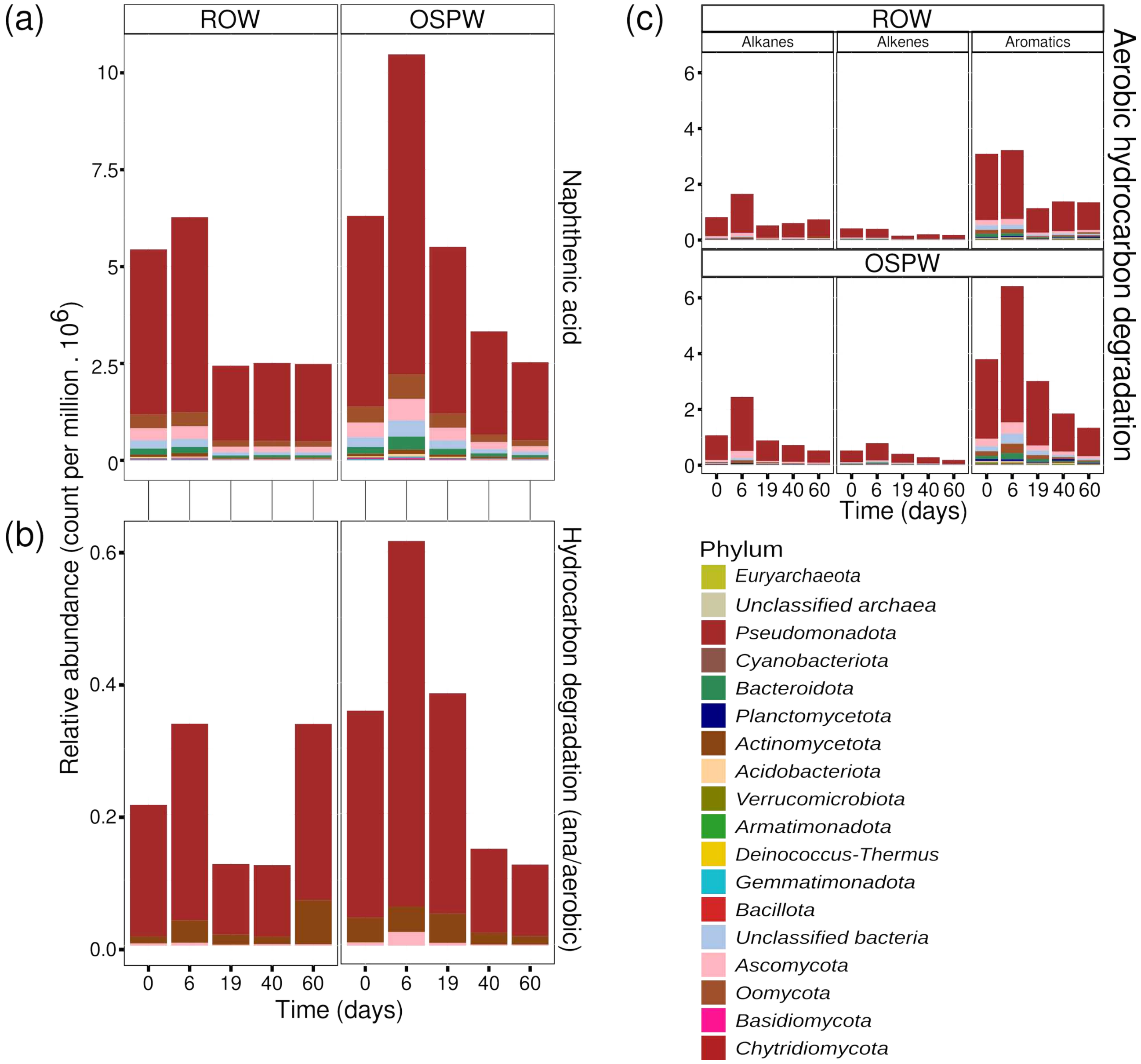


***Fig. S3. Abundance and phylum-level taxonomic origin of expressed genes potentially involved in naphthenic acid degradation.*** *The figure shows the average gene abundance at the phylum level, expressed as counts per million (CPM). (a) Genes previously associated with naphthenic acid degradation, identified using a custom Hidden Markov Model (HMM) database. (b) Aerobic and anaerobic hydrocarbon degradation marker genes identified using the CANT-HYD database. (c) Aerobic hydrocarbon degradation genes profiled using the HADEG database. The data were obtained from the roots of Typha latifolia grown in mesocosms mimicking constructed wetland systems irrigated with either reverse osmosis water (ROW) or oil sands process-affected water (OSPW) at five time points: 0, 6, 19, 40, and 60. Time 0 is before addition of OSPW; 6-60 represent sampling days after adding OSPW or refilling with ROW.*

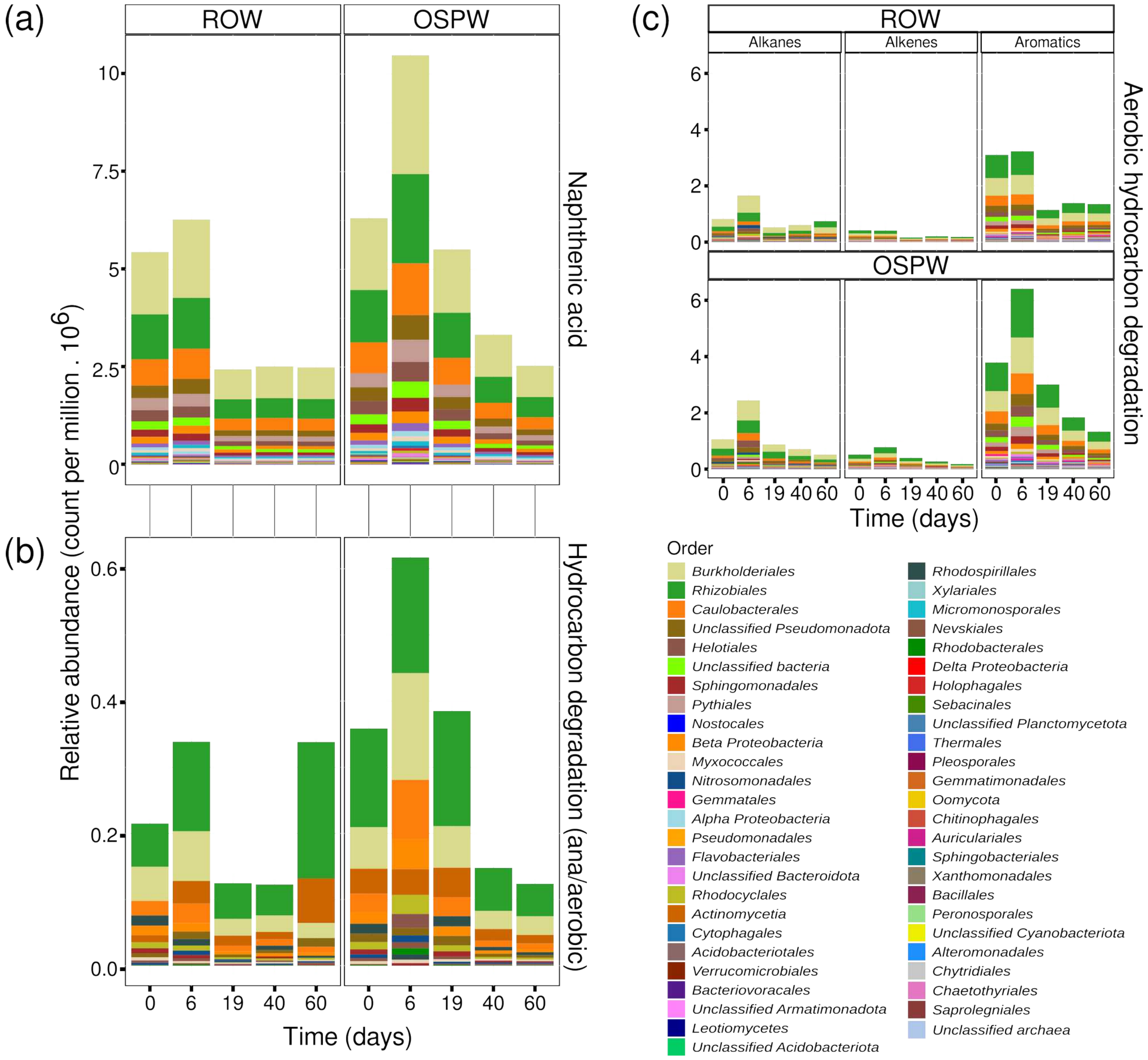


***Fig. S4. Abundance and order-level taxonomic origin of expressed genes potentially involved in naphthenic acid degradation.*** *The figure shows the average gene abundance at the order level, expressed as counts per million (CPM). (a) Genes previously associated with naphthenic acid degradation, identified using a custom Hidden Markov Model (HMM) database. (b) Aerobic and anaerobic hydrocarbon degradation marker genes identified using the CANT-HYD database. (c) Aerobic hydrocarbon degradation genes profiled using the HADEG database. The data were obtained from the roots of Typha latifolia grown in mesocosms mimicking constructed wetland systems irrigated with either reverse osmosis water (ROW) or oil sands process-affected water (OSPW) at five time points: 0, 6, 19, 40, and 60. Time 0 is before addition of OSPW; 6-60 represent sampling days after adding OSPW or refilling with ROW.*

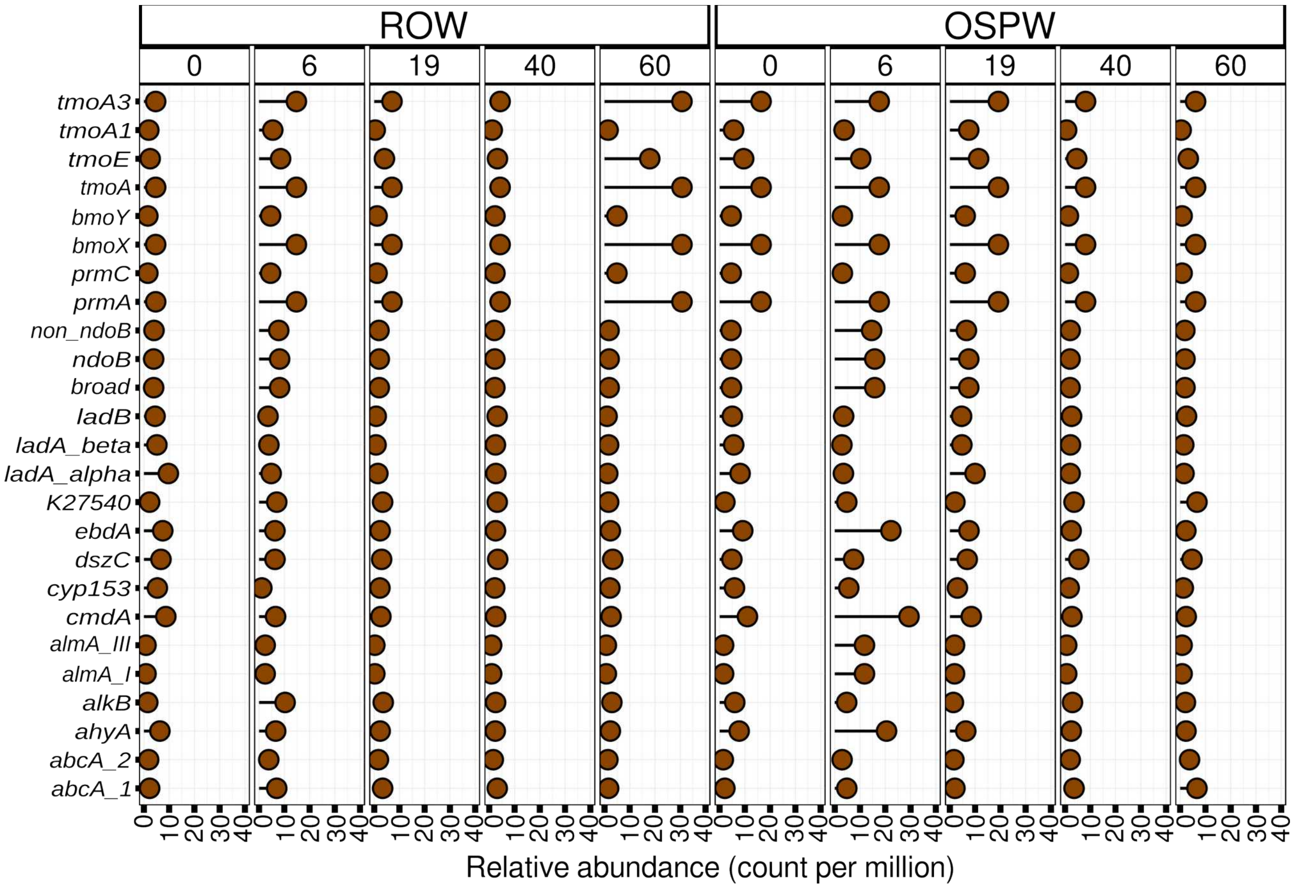


***Fig. S5. Abundance aerobic and anaerobic hydrocarbon degradation marker genes identified using the CANT-HYD database.*** *The figure shows the average gene abundance expressed as counts per million (CPM). The data were obtained from the roots of Typha latifolia grown in mesocosms mimicking constructed wetland systems irrigated with either reverse osmosis water (ROW) or oil sands process-affected water (OSPW) at five time points: 0, 6, 19, 40, and 60. Time 0 is before addition of OSPW; 6-60 represent sampling days after adding OSPW or refilling with ROW.*

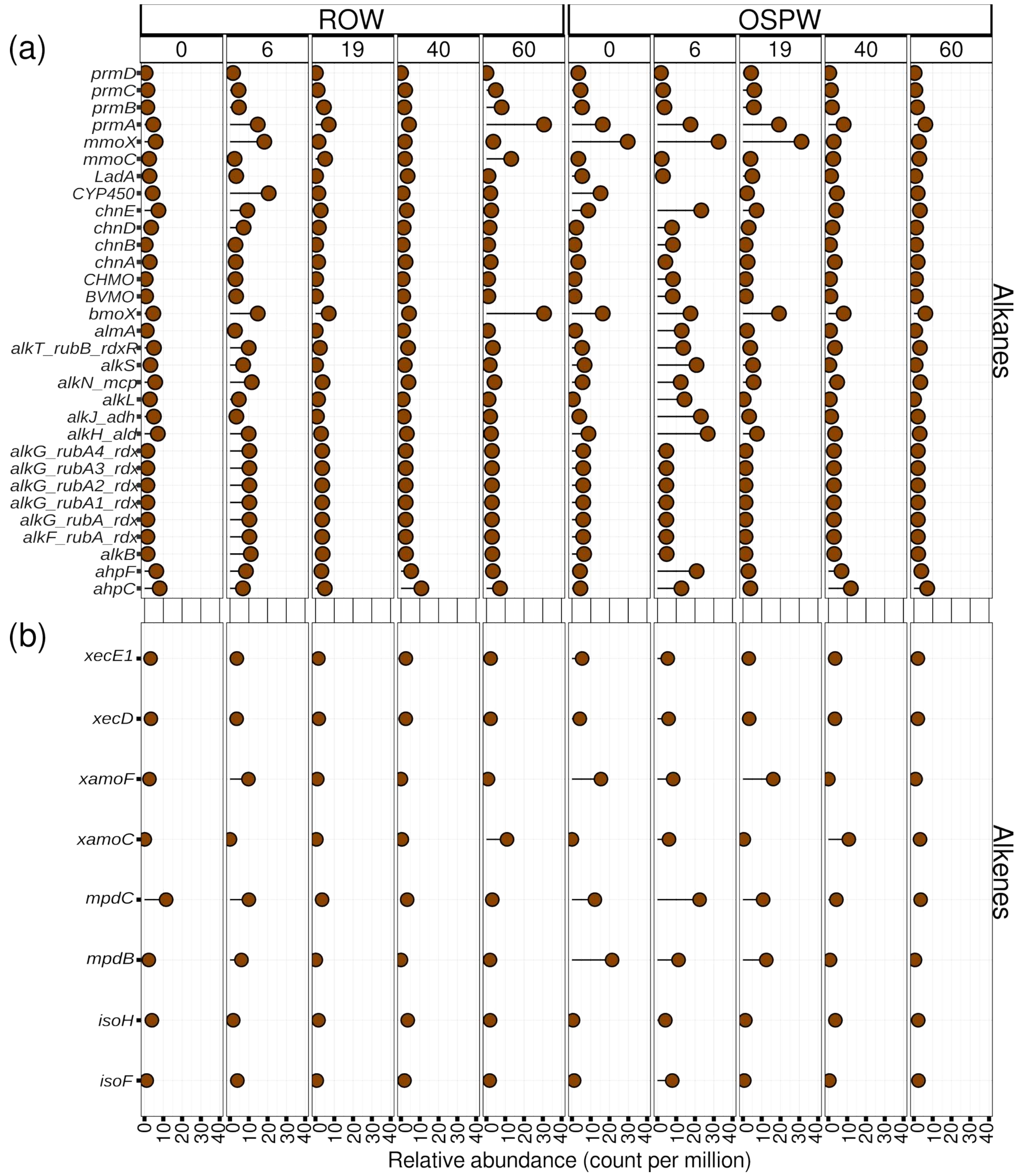


***Fig. S6. Abundance aerobic hydrocarbon degradation marker genes identified using the HADEG database.*** *The figure shows the average gene abundance expressed as counts per million (CPM). The genes for degradation of (a) alkanes and (b) alkenes. The genes were profiled using a curated database of hydrocarbon aerobic degradation enzymes and genes (HADEG) database containing proteins and genes involved in alkane, alkene and aromatic aerobic degradation. The data were obtained from the roots of Typha latifolia grown in mesocosms mimicking constructed wetland systems irrigated with either reverse osmosis water (ROW) or oil sands process-affected water (OSPW) at five time points: 0, 6, 19, 40, and 60. Time 0 is before addition of OSPW; 6-60 represent sampling days after adding OSPW or refilling with ROW.*

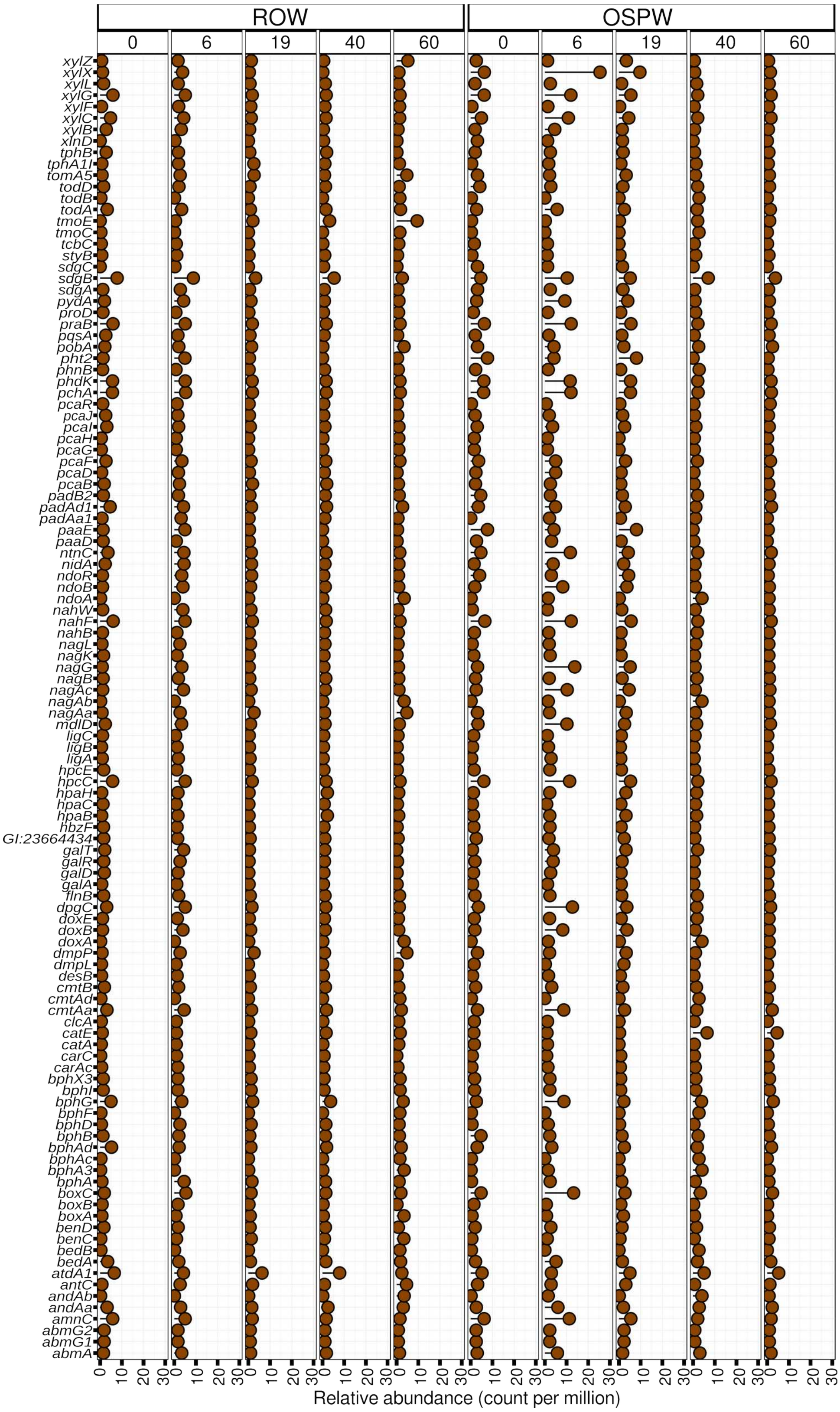


***Fig. S6c. Abundance aerobic aromatic hydrocarbon degradation marker genes identified using the HADEG database.*** *The figure shows the average gene abundance expressed as counts per million (CPM). Time 0 is before addition of OSPW; 6-60 represent sampling days after adding OSPW or refilling with ROW.*

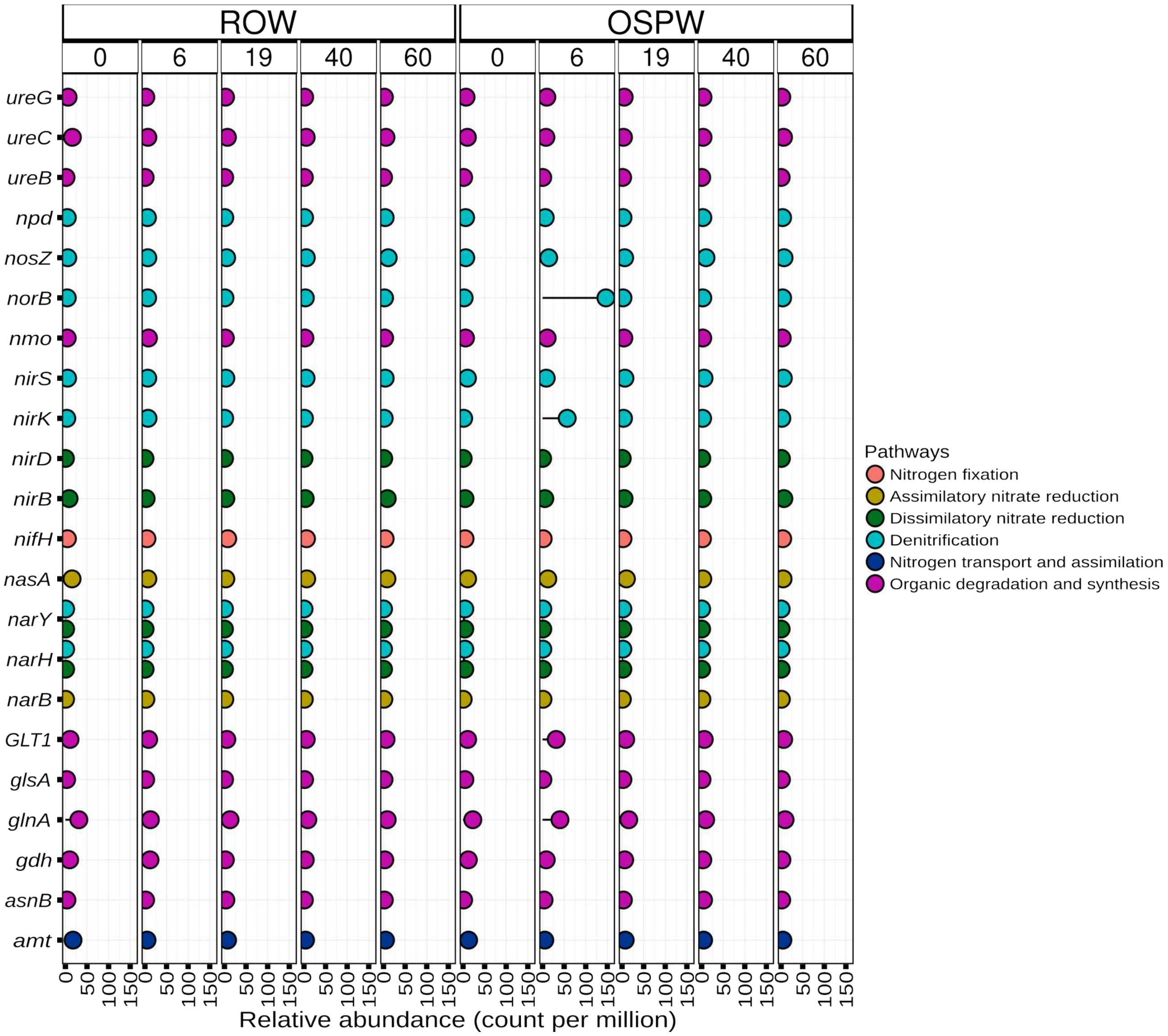


***Fig. S7. Abundance of genes involved in nitrogen metabolism using the NCyc database.*** *The figure shows the average gene abundance expressed as counts per million (CPM). The genes were profiled using a curated integrative database for fast and accurate metagenomic profiling of nitrogen cycling genes (NCyc) containing 68 gene (sub)families and covers eight N cycle processes. The data were obtained from the roots of Typha latifolia grown in mesocosms mimicking constructed wetland systems irrigated with either reverse osmosis water (ROW) or oil sands process-affected water (OSPW) at five time points: 0, 6, 19, 40, and 60. Time 0 is before addition of OSPW; 6-60 represent sampling days after adding OSPW or refilling with ROW.*

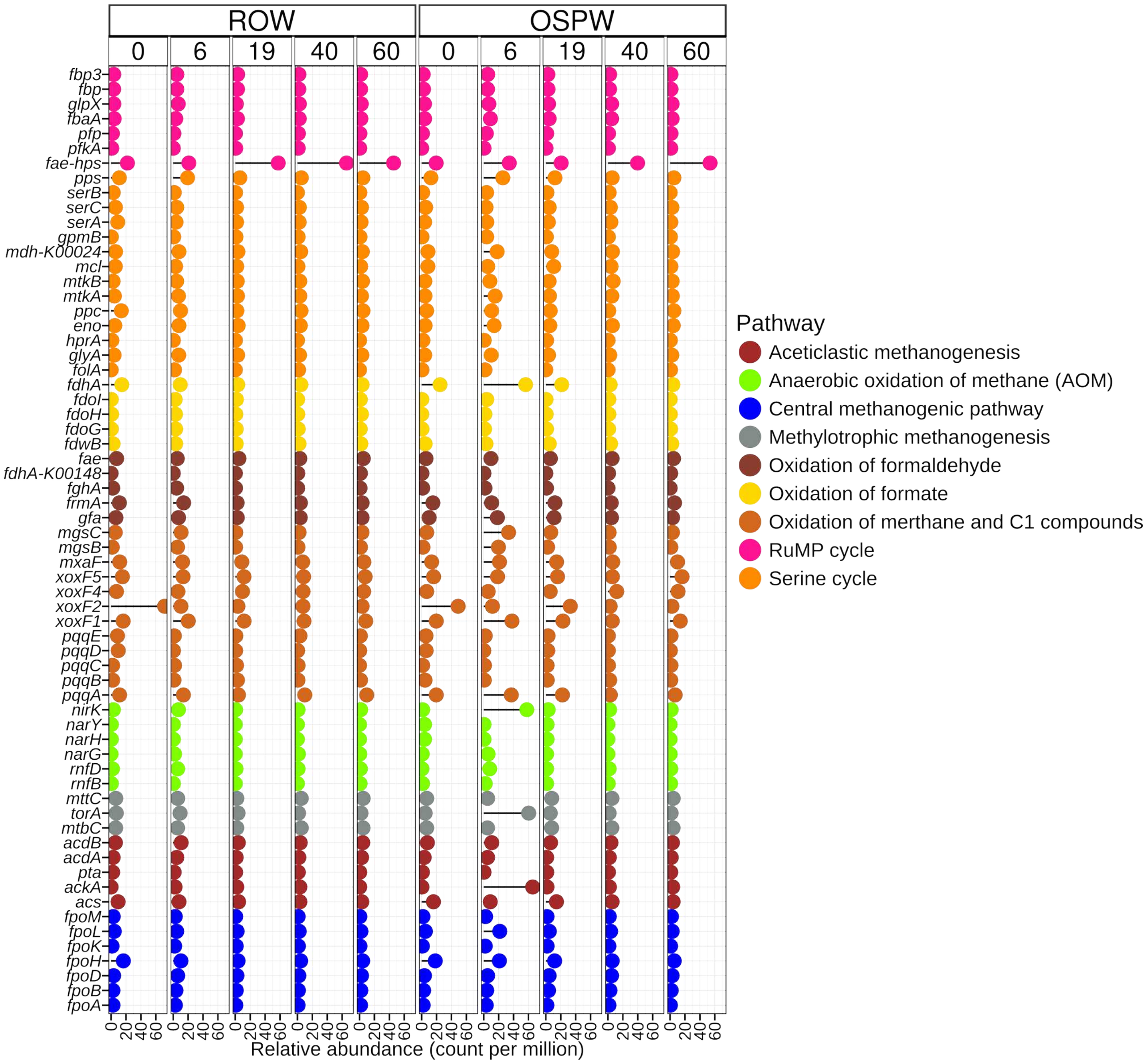


***Fig. S8. Abundance of genes involved in sulfur metabolism using the SCycDB database.*** *The figure shows the average gene abundance expressed as counts per million (CPM). The genes were profiled using a curated functional gene database for metagenomic profiling of sulfur cycling pathways (SCycDB) containing 207 gene families. The data were obtained from the roots of Typha latifolia grown in mesocosms mimicking constructed wetland systems irrigated with either reverse osmosis water (ROW) or oil sands process-affected water (OSPW) at five time points: 0, 6, 19, 40, and 60. Time 0 is before addition of OSPW; 6-60 represent sampling days after adding OSPW or refilling with ROW.*

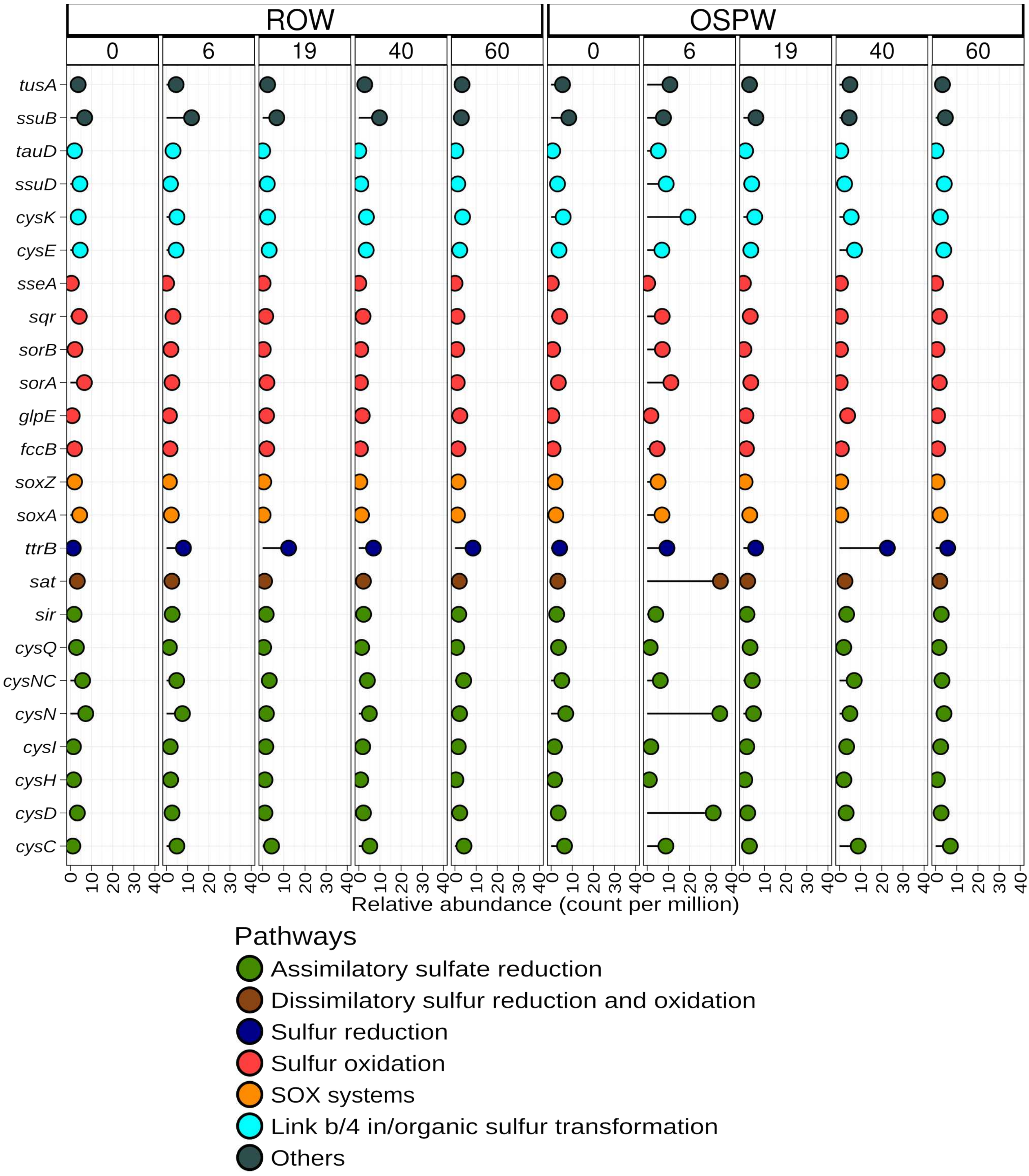


***Fig. S9. Abundance of genes involved in phosphorus metabolism using the PCyCDB database.*** *The figure shows the average gene abundance expressed as counts per million (CPM). The genes were profiled using a comprehensive and accurate database for fast analysis of phosphorus cycling genes (PCyCDB) containing 139 gene families and 10 P metabolic processes. Time 0 is before addition of OSPW; 6-60 represent sampling days after adding OSPW or refilling with ROW.*

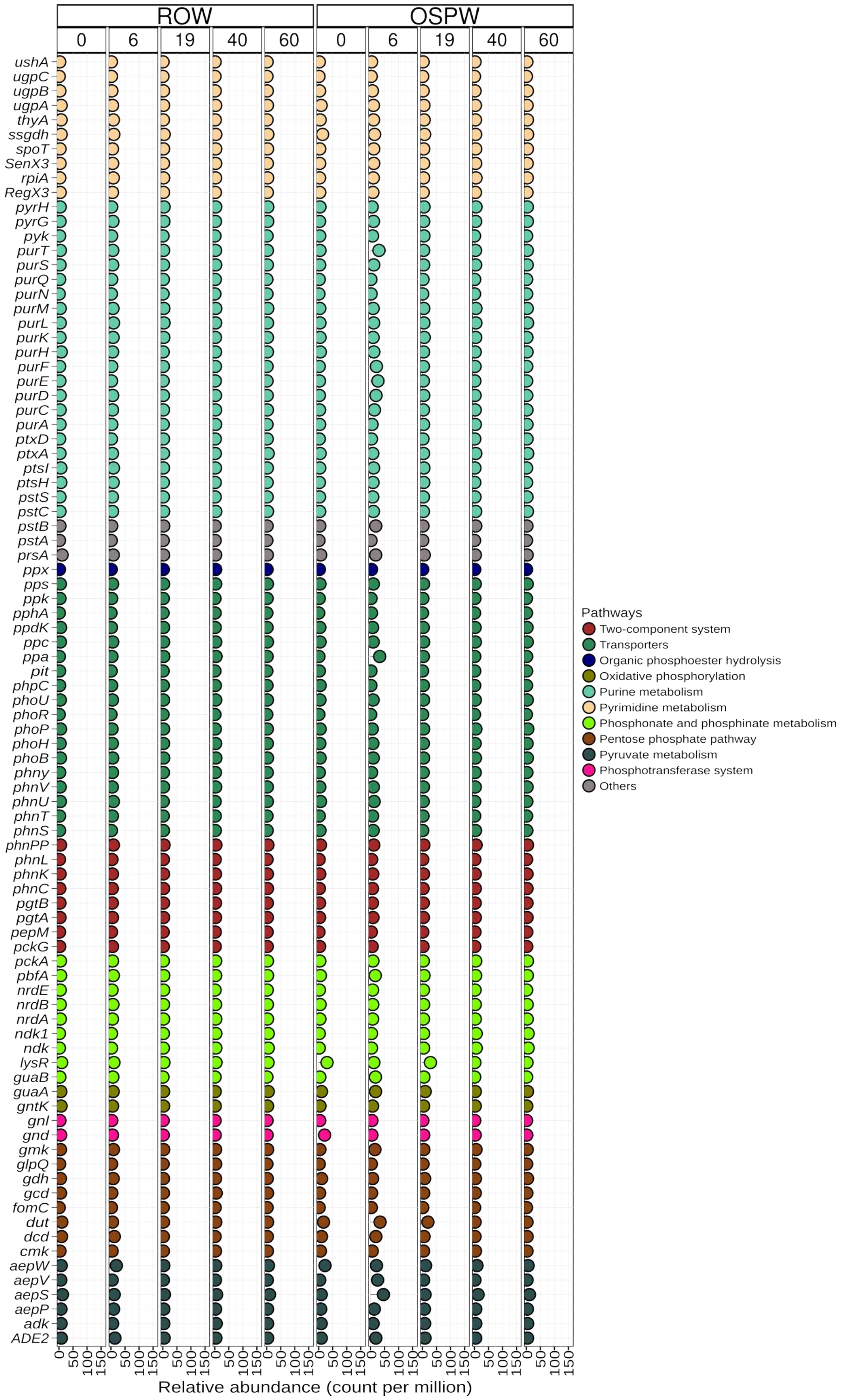


***Fig. S10. Abundance of genes involved in methane metabolism using the MCycDB database.*** *The figure shows the average gene abundance expressed as counts per million (CPM). The genes were profiled using a curated database for comprehensively profiling methane cycling processes of environmental microbiomes (MCycDB) containing 298 methane cycling gene families covering 10 methane metabolism pathways. The data were obtained from the roots of Typha latifolia grown in mesocosms mimicking constructed wetland systems irrigated with either reverse osmosis water (ROW) or oil sands process-affected water (OSPW) at five time points: 0, 6, 19, 40, and 60. Time 0 is before addition of OSPW; 6-60 represent sampling days after adding OSPW or refilling with ROW.*

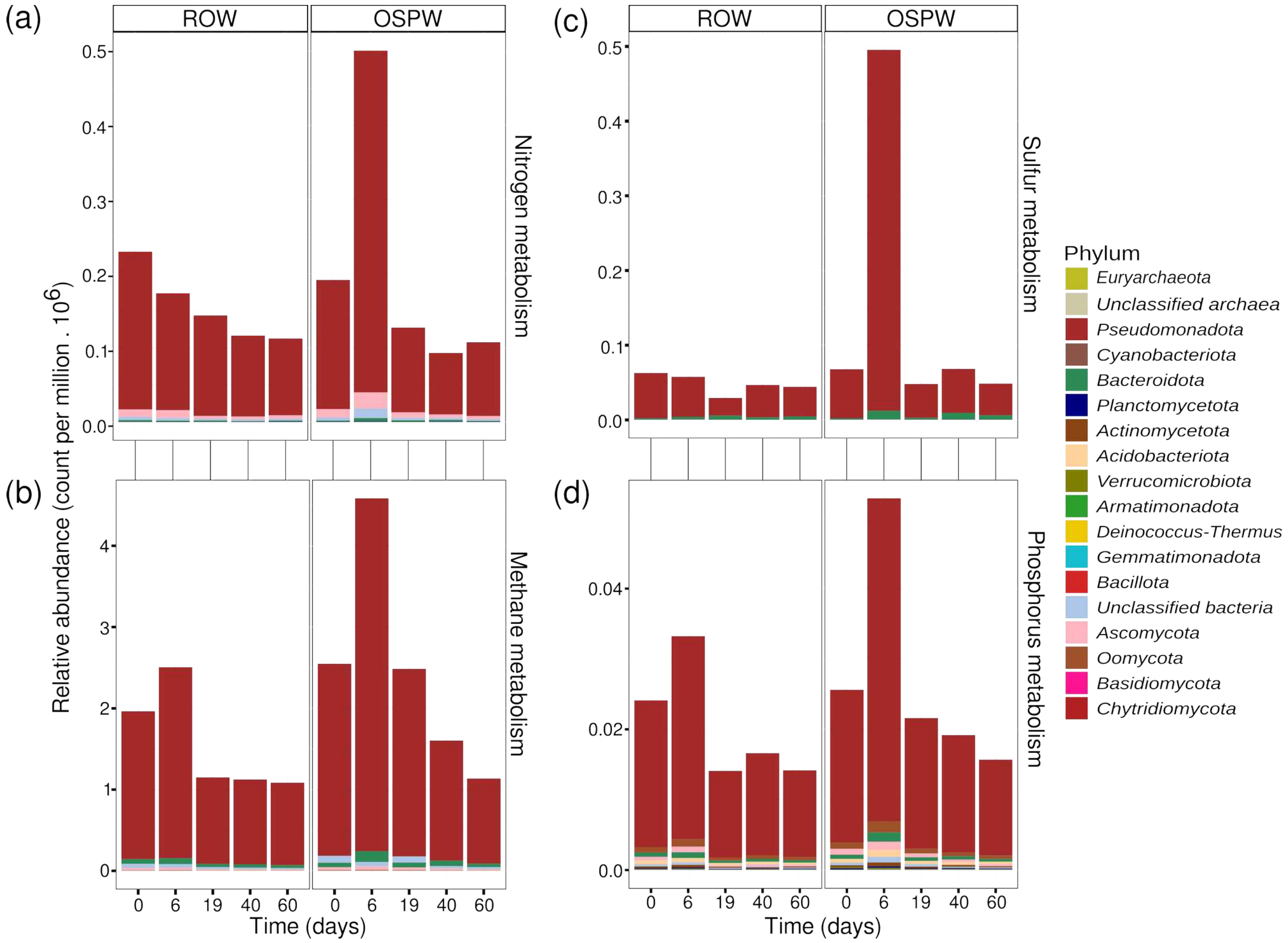


***Fig. S11. Abundance and phylum-level taxonomic origins of genes involved in nutrient cycling.*** *The figure shows the average gene abundance expressed as counts per million (CPM) and summed at the phylum level. The genes for degradation of (a) nitrogen, (b) methane, (c) sulfur and (d) phosphorus metabolism. The data were obtained from the roots of Typha latifolia grown in mesocosms mimicking constructed wetland systems irrigated with either reverse osmosis water (ROW) or oil sands process-affected water (OSPW) at five time points: 0, 6, 19, 40, and 60. Time 0 is before addition of OSPW; 6-60 represent sampling days after adding OSPW or refilling with ROW.*

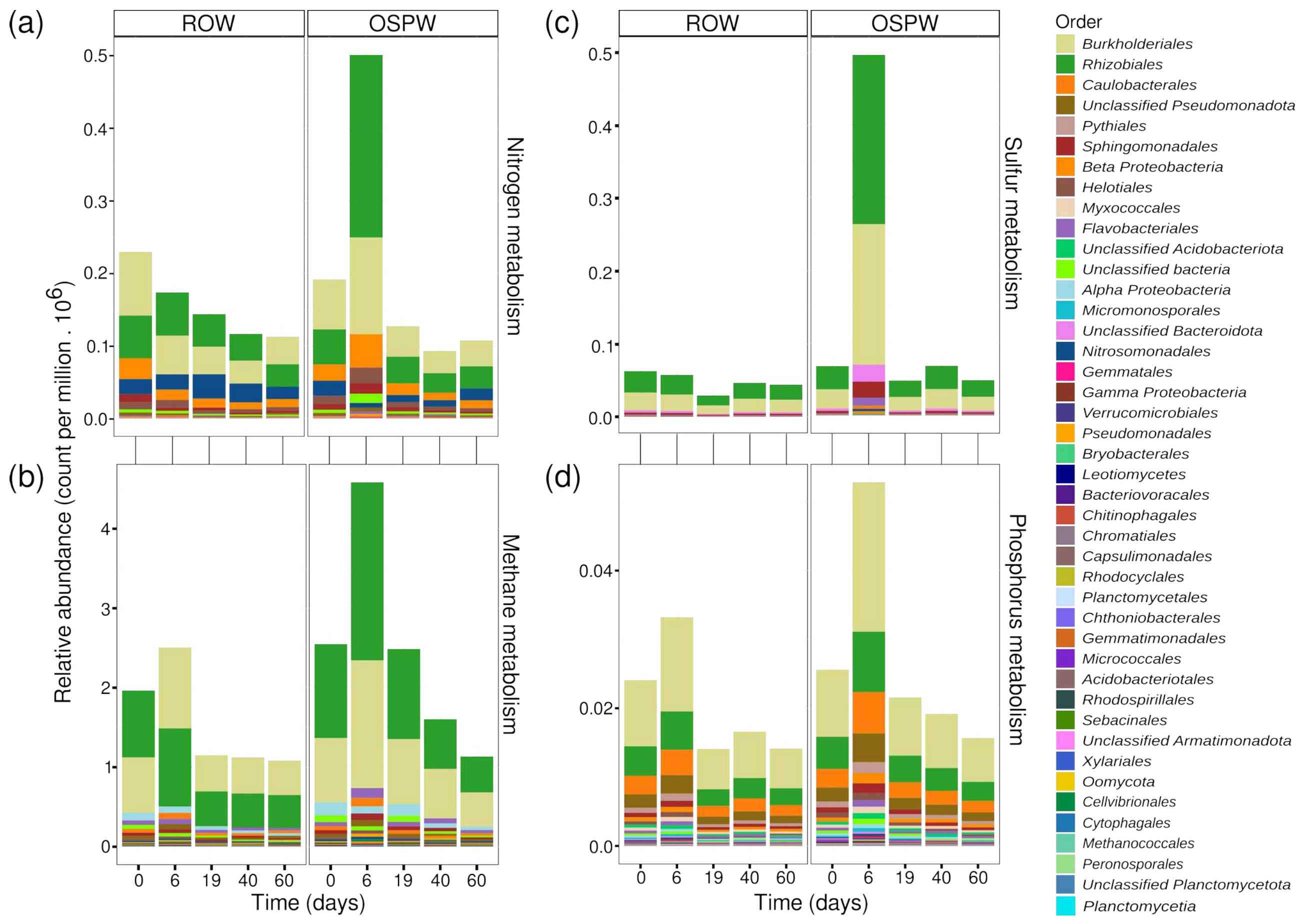


***Fig. S11. Abundance and order-level taxonomic origins of genes involved in nutrient cycling.*** *The figure shows the average gene abundance expressed as counts per million (CPM) and summed at the order level. The genes for degradation of (a) nitrogen, (b) methane, (c) sulfur and (d) phosphorus metabolism. The data were obtained from the roots of Typha latifolia grown in mesocosms mimicking constructed wetland systems irrigated with either reverse osmosis water (ROW) or oil sands process-affected water (OSPW) at five time points: 0, 6, 19, 40, and 60. Time 0 is before addition of OSPW; 6-60 represent sampling days after adding OSPW or refilling with ROW.*
